# Drebrin Regulates Collateral Axon Branching in Cortical Layer II/III Somatosensory Neurons

**DOI:** 10.1101/2023.06.21.545958

**Authors:** Joelle M. Dorskind, Sriram Sudarsanam, Randal A. Hand, Jakub Ziak, Maame Amoah-Dankwah, Luis Guzman-Clavel, John Lee Soto-Vargas, Alex L. Kolodkin

**Affiliations:** Solomon H. Snyder Department of Neuroscience, Johns Hopkins Kavli Neuroscience Discovery Institute, The Johns Hopkins University School of Medicine, Baltimore, MD, 21205

**Keywords:** Lamination, Circuitry, Cytoskeleton, Axon, Branching, Microtubules, Actin, Drebrin

## Abstract

Proper cortical lamination is essential for cognition, learning, and memory. Within the somatosensory cortex, pyramidal excitatory neurons elaborate axon collateral branches in a laminar-specific manner that dictates synaptic partners and overall circuit organization. Here, we leverage mouse models, single-cell labeling and imaging approaches to identify intrinsic regulators of laminar-specific collateral, also termed interstitial, axon branching. We developed new approaches for the robust, sparse, labeling of layer II/III pyramidal neurons to obtain single-cell quantitative assessment of axon branch morphologies. We combined these approaches with cell-autonomous loss-of-function (LOF) and over-expression (OE) manipulations in an *in vivo* candidate screen to identify regulators of cortical neuron axon branch lamination. We identify a role for the cytoskeletal binding protein drebrin (Dbn1) in regulating layer II/III cortical projection neuron (CPN) collateral axon branching *in vitro.* LOF experiments show that Dbn1 is necessary to suppress the elongation of layer II/III CPN collateral axon branches within layer IV, where axon branching by layer II/III CPNs is normally absent. Conversely, *Dbn1* OE produces excess short axonal protrusions reminiscent of nascent axon collaterals that fail to elongate. Structure-function analyses implicate Dbn1^S142^ phosphorylation and Dbn1 protein domains known to mediate F-actin bundling and microtubule (MT) coupling as necessary for collateral branch initiation upon Dbn1 OE. Taken together, these results contribute to our understanding of the molecular mechanisms that regulate collateral axon branching in excitatory CPNs, a key process in the elaboration of neocortical circuit formation.

**Significance Statement:** Laminar-specific axon targeting is essential for cortical circuit formation. Here, we show that the cytoskeletal protein drebrin (Dbn1) regulates excitatory layer II/III cortical projection neuron (CPN) collateral axon branching, lending insight into the molecular mechanisms that underlie neocortical laminar-specific innervation. To identify branching patterns of single cortical neurons *in vivo*, we have developed tools that allow us to obtain detailed images of individual CPN morphologies throughout postnatal development and to manipulate gene expression in these same neurons. Our results showing that Dbn1 regulates CPN interstitial axon branching both *in vivo* and *in vitro* and may aid in our understanding of how aberrant cortical neuron morphology contributes to dysfunctions observed in Autism Spectrum Disorder (ASD) and epilepsy.

## Introduction

The establishment of complex neural circuits requires spatial and temporal coordination of neuronal responses to guidance cues. Growing axons and dendrites must filter a myriad of signals to identify relevant cues and navigate to their appropriate targets during neocortical development (Sanes and Yamagata, 2009; Kolodkin and Tessier-Lavigne, 2011). The primary somatosensory cortex (S1) is organized into six molecular layers, each of which contains distinct cell populations (Greig et al., 2013). Neurons within each cortical layer extend axons over long distances to establish connections with cortical or subcortical regions, providing a scaffold for global cortical circuit organization (Molyneaux et al., 2007). Excitatory neurons within each layer form many of their local synaptic connections through the elaboration of interstitial (also termed “collateral”) axon branches that extend from the primary axon shaft in a laminar-specific fashion (Molyneaux et al., 2007; Kalil and Dent, 2014; Hand et al., 2015).

Excitatory layer II/III cortical projection neurons (CPNs) have unique and stereotypic projection and branching patterns and are thus a robust model for studying the molecular mechanisms underlying cortical circuit formation and organization. For example, layer II/III CPNs generate collateral branches mainly in S1 layers II/III and V, and seldom in layers IV and VI, both in the ipsilateral and contralateral brain hemispheres (Borrell and Callaway, 2002; Hand et al., 2015). Studies utilizing layer II/III CPNs reveal several molecular mechanisms underlying cortical neuron polarization, migration, and axon extension (Barnes et al., 2007; Hand and Polleux, 2011; Courchet et al., 2013; Martinez-Garay et al., 2016; Zhang et al., 2018). However, a comprehensive view of the molecules that regulate laminar-specific neuronal process elaboration via collateral axon branching of projection neurons within the cortex is still lacking (Shepherd and Rowe, 2017; Dorskind and Kolodkin, 2021). In this study, we use layer II/III neurons to investigate molecular regulators of collateral axon branching *in vivo*.

Cytoskeletal rearrangements are critical for regulating neuronal morphology during development (Dent et al., 2003; Kapitein and Hoogenraad, 2015). Previous work describes mechanisms and cytoskeletal regulators that are needed to form collateral axon branches *in vitro* (Dent et al., 2003; Kalil and Dent, 2014), including the promotion of microtubule (MT) physical engagement with the actin meshwork, which serves as a precursor for the formation of collateral axon branches (Gibson and Ma, 2011; Hand et al., 2015; Ketschek et al., 2016). Drebrin1 (Dbn1) is a cytoskeletal regulatory protein that contains both MT and actin binding sites. Drebrin binds to parallel actin filaments at the base of filopodia and interacts with End Binding Protein 3 (EB3) located at the plus ends of MTs (Worth et al., 2013; Ketschek et al., 2016; Gordon-Weeks, 2017).

There are two major mammalian drebrin isoforms, E and A, and they are produced by alternative splicing from a single *Dbn1* transcript. Dbn1-E (referred to here and below as “Dbn1”) plays numerous roles in neurite outgrowth and branching (Shirao et al., 2017). Dbn1 over-expression in dorsal root ganglion (DRGs) neurons *in vivo* and *in vitro* increases filapodial-like protrusions along axons (Spillane et al., 2012; Ketschek et al., 2016) and increases the number of collateral axon branches along the midbrain medial longitudinal fasciculus (MLF) (Dun et al., 2012). Conversely, *Dbn1* knockdown limits axon growth and branching *in vitro* in DRG and embryonic cortical neurons, respectively (Ketschek et al., 2016; Poobalasingam et al., 2022).

These studies demonstrate that Dbn1 figures prominently in neurite outgrowth and axon branching and suggest that Dbn1 is an attractive candidate for contributing to laminar-specific innervation through its axon collateral branch initiation and stabilization functions. However, an *in vivo* role for Dbn1 in regulating excitatory CPN interstitial axon branching has yet to be determined. Here, we show at the single cell level that Dbn1 is necessary for the suppression of collateral axon branching by layer II/III somatosensory CPNs *in vivo* and *in vitro*. These observations provide insight into mechanisms necessary for establishing axon innervation patterns during neocortical development.

### Materials and Methods

#### Animals

*Dbn1^flox^* mice were a gift from Dr. Jonathan Sitber (Duke) (Stiber et al., 2016). Timed pregnant embryonic day 15.5 CD1 females were obtained from Charles River Laboratory (Strain code 022). The day of birth was designated as P0. Mice of both sexes were used in all experiments and were housed in a 12:12 light-dark (LD) cycle. This research was carried out in strict accordance with the recommendations in the Guide for the Care and Use of Laboratory Animals of the NIH. The animal protocol was approved by the Animal Care and Use Committees of the Johns Hopkins University School of Medicine.

#### Immunohistochemistry

Mice were deeply anesthetized and perfused with ice-cold 1X PHEM (pH 6.9) followed by 4% paraformaldehyde in 1X PHEM with 10% sucrose and 0.1% Triton X-100 (4%PFA/PHEM). 4X PHEM solution contained: 60mM PIPES (Amresco (0169-250G), 25mM HEPES (Sigma (H3375-500G)), 5mM EGTA (Amresco (0732-1006)), 1mM MgCl2 (Sigma (M8266-100G)) and ddH2O to 1L. Brains were promptly dissected and post-fixed in 4% PFA/PHEM, PBS (pH 7.4) for 2 hrs at room temperature. Fixed brains were washed in PBS and 250-μm coronal sections were prepared using a vibratome. Brain sections were permeabilized and blocked in permeabilization buffer (3% BSA, 0.3% Triton X-100, and PBS [pH 7.4]) with rocking overnight at 4°C. Primary antibodies were diluted in permeabilization buffer and secondary antibodies and DAPI were diluted in permeabilization buffer with 10% goat serum. The brain sections were incubated with antibody solutions overnight rocking at 4°C. After antibody incubation, brain sections were washed in PBS (pH 7.4) five times for 30 minutes with shaking at room temperature. Prior to imaging, floating brain sections were mounted onto glass slides and dried. Fluoro Gel with DABCO (Electron Microscopy Sciences) was applied to each slide and a coverslip mounted on top. Primary antibodies used were: chicken anti-GFP (Aves, GFP-1020, 1:1000), rabbit anti-dsRed (Takara, 632496, 1:1000), which was used to detect mCherry signal, and mouse anti-Drebrin (Mbl,1:500). Cell nuclei were stained with 4’,6-Diamidino-2-Phenylindole (DAPI, 1:1000, ThermoFisher D1306).

#### Targeted *in utero* electroporation

Methods were adapted from (Hand et al., 2015). Timed pregnant embryonic day 15.5 CD1 or *Dbn1^Fl/+^* females were deeply anesthetized with either 2,2,2 tribromoethanol 2.5% in PBS (pH 7.4) or isoflurane. The laparotomy site was shaved, cleaned, a small longitudinal incision (1.5–2 cm) was made, and the embryos removed and rinsed with sterile PBS. The lateral ventricles were injected with small volumes of DNA solutions in PBS (pH 7.4) with fast green dye. Embryos were electroplated with gene paddles (Harvard Apparatus) using a BTX square pulse electroporator: 30–40 V, 3 Å∼ 50-ms pulses with a 950-ms interval. The pain control included subcutaneous injection of local anesthetic (Bupivacaine) prior to skin incision and subcutaneous injection of Buprenorphine after surgery. All procedures were conducted in accordance to IUCAC-approved protocols.

The following combination of plasmids were used for each electroporation approach described in this manuscript (details for below):

1. Single Recombinase Approach: *pCAG-Cre (or pCAG-CreERT2)* (0.5μg/ul), *pCAG-LSL-mCherry* (1μg/ul), *pCAG-*GENEX*-IRES-GFP* (2μg/ul),,
2. Bipartite Approach: *pCAG-CAG-CreERT2* (0.5μg/ul)*, pEF1-Flex-FlpO* (0.5μg/ul*,, pCAG-FSF-mCherry* (1μg/ul), *pCAG-FSF-*GENEX*-IRES-GFP* (or tagged YFP) (2μg/ul),
3. Modified Bipartite Approach: *pCAG-CreERT2* (0.3-0.5μg/ul), *pAAV-TRE-DIO-FlpO* (0.01-0.015μg/ul),, *pCAG-FSF-turboRFP-IRES-tTA* (0.75-1μg/ul),, *pCAG-FSF-Dbn1S142D-YFP* (1.5μg/ul)

#### Plasmids

For cloning of all over-expression constructs (including DN and CA constructs), primers were designed with the addition of restriction enzyme sites to clone into our *pCIG2* (pCAG-IRES-GFP) plasmid to amplify gene of interest from P5 mouse cDNA. CA-GSK3β was modified from HA GSK3 beta S9A pcDNA3 a gift from Jim Woodgett (Addgene plasmid # 14754; http://n2t.net/addgene:14754; RRID:Addgene_14754). *pCAG-YEPT* was subcloned from pYpet-C1, which was a gift from Klaus Hahn (Addgene plasmid # 22780; http://n2t.net/addgene:22780; RRID:Addgene_22780). *pCAG-CreERT2* was a gift from Connie Cepko (Addgene plasmid # 13777; http://n2t.net/addgene:13777; RRID:Addgene_13777). *pCAG-FlpO* was a gift from Massimo Scanziani (Addgene plasmid # 60662; http://n2t.net/addgene:60662; RRID:Addgene_60662). This plasmid was then modified to create *pCAG-FlpO*, which was cloned from a *pEF1-Flex* construct. *pEF1-Flex* was generated from the *pEF1-EGFP* backbone by replacing a Flex multiple cloning site cassette with EGFP. For plasmids containing *pCAG-FSF*, the stop cassette was amplified from *pCAG-LSL-4* plasmid with primers encoding the FRT sites and clones into *pCAG-LSL* with the restriction sites PacI/XhoI. For *pCAG-mCherry* (used for *pCAG-FSF-mCherry*), mCherry was PCR amplified from Cortactin-pmCherry-C1 and cloned into pCAG-LSL using EcoRI/NotI restriction sites. Cortactin-pmCherryC1 was a gift from Christien Merrifield (Addgene plasmid # 27676; http://n2t.net/addgene:27676; RRID:Addgene_27676). Dbn1 phospho-mutant constructs were modified from gifts from Phillip Gordon-Weeks: Dbn1-YFP (Addgene plasmid # 40359; http://n2t.net/addgene:40359; RRID:Addgene_40359), Dbn1-S142D-YFP (Addgene plasmid # 58336; http://n2t.net/addgene:58336; RRID:Addgene_58336), and Dbn1-S142A-YFP (Addgene plasmid # 58335; http://n2t.net/addgene:58335; RRID:Addgene_58335). Dbn1-YFP in each of these constructs was then subcloned into *pCAG-FSF-IRES-GFP*. For deletion constructs, regions of Dbn1 were amplified using PCR with primers containing XHO1/NHE1 restriction sites. These fragments were then cloned into *pCAG-FSF-IRES-GFP*.

For general labeling of neurons with eGFP or mCherry, a pCAG-eGFP plasmid at a concentration of 2 μg/μl or a pCAG-mCherry plasmid at a concentration of 1 μg/μl were used. For constitutive sparse labeling with eGFP (*pCAG-LSL-eGFP,* 2 μg/μl) or mCherry (*pCAG-LSL-mCherry*,1 μg/μl) and Cre (*pCAG-Cre* at concentrations ranging between 1-10 ng/μl were co-electroporated single recombinase screening approach). For conditional, post-migration sparse labeling using the bipartite approach with eGFP (*pCAG-FSF-eGFP*, 2 μg/μl) or DNA construct of interest (*pCAG-FSF-*GENEX, 2 μg/μl), mCherry (*pCAG-FSF-mCherry* at, 1 μg/μl), CreERT2 (*pCAG_CreERT2*, 500 ng/μl), and Cre-dependent FlpO (pEF1-Flex-FlpO, 500 ng/μl were co-electroporated. In cases where a conditional mouse line was used, mCherry (*pCAG-FSF-mCherry*, 1 μg/μl), CreERT2 (*pCAG_CreERT2*, 500ng/μl), and Cre-dependent FlpO (*pEF1-Flex-FlpO*, 500 ng/μl) were co-electroporated.

For Dbn1 structure-function analyses, domain deletion-specific plasmids were generated based on published sequences and strategies (Shirao and Sekino, 2017). *Dbn1ΔADF-H* (residues 8-134 were deleted), *Dbn1ΔAB1/2* included both the CC and Hel domains (residues 173-317 were deleted), *Dbn1ΔPP* (residues 410-419 were deleted), *Dbn1ΔN-terminal* (residues 1-317 were deleted), *Dbn1ΔC-Terminal* (residues 365-661 were deleted).

For Dbn1 rescue experiments using the *Dbn1^flox^* allele the following constructs were used in lieu of the original bipartite constructs (Luo et al., 2016; Lin et al., 2018): *pAAV-TRE-DIO-FlpO* was a gift from Minmin Luo (Addgene plasmid # 118027; http://n2t.net/addgene:118027; RRID:Addgene_118027), *pCAG-FSF-turboRFP-IRES-tTA* was a gift from Takuji Iwasato (Addgene plasmid # 85038; http://n2t.net/addgene:85038; RRID:Addgene_85038).

#### Tamoxifen injections

This protocol was adapted from (Pitulescu et al., 2010). Tamoxifen solution was prepared by dissolving tamoxifen powder (Sigma-Aldrich, T5648) in 100% ethanol to a concentration of 10 mg/ml. Corn oil (MP Biomedicals, 901414) was added to make a working concentration of 1 mg/ml. For two consecutive days (P1 + P2), 30μl of this solution was administered via intragastric injection directly into a visible milk spot.

#### *Ex vivo* electroporation and primary neuron cell culture

Primary mouse cortical neurons were derived from E15.5 wild-type or indicated mutant embryos. Electroporation of E15.5 cortices was performed by injecting small volumes of DNA solutions in PBS (pH 7.4) with fast green dye into the lateral ventricles of isolated embryonic mouse heads that were decapitated and placed in complete HBSS. The lateral ventricles were injected with approximately 1μl of DNA solutions in PBS (pH 7.4) with fast green dye, until the ventricles were just full. Embryos were electroplated with gene paddles (Harvard Apparatus) using a BTX square pulse electroporator: 30–40 V, 3 Å∼ 50-ms pulses with a 950-ms interval. The positive electrode was placed above the differentiating areas of the primary somatosensory cortex. Primary mouse cortical neurons were derived from E15.5 wild-type or indicated mutant embryos. The following protocol was modified and adapted from (Polleux and Ghosh, 2002).

Cortical tissues were dissected from embryos of both sexes and digested in a dissociation medium (DM) containing 98mM Na2SO4, 30mM K2SO4, 5.8mM MgCl2, 0.25mM CaCl2,1mM Hepes (pH 7.4), 20mM glucose, 0.001% Phenol red, and 0.124mN NaOH. 400 units of papain (Worthington, LS003126) and 6.4mg Cysteine were added to the DM for a 20mL total solution. Solution was mixed and pH was adjusted with 0.1 N NaOH to ∼7.4 monitored by color of solution (pink, too basic; yellow, too acidic). The solution was then filtered through a 0.2μm syringe filter. Cortices were incubated in solution for approximately 15 min in a 37℃ water bath, or until cortices were “stringy” in appearance. Cortices were then placed into a heavy inhibitory (HI) solution containing 6ml of DM and 60mg BSA (pH 7.4), warmed to 37℃, and then transferred to a light. inhibitory (LI) solution containing 9ml of DM and 1ml of HI. Cortices were then washed three times in Neurobasal Medium containing 5% Penicillin (10,000 units/ml)-streptomycin (10mg/ml), and then underwent mechanical trituration using a 1ml pipette tip. Cells were counted and plated at a density of 30-70,000 cells/well in a Serum-Free Medium on pre-treated glass coverslips in either 12 or 24 well Nunc plates. A 50ml total solution was made containing Neurobasal medium, L-glutamine (200mM) 250ul, penicillin (10,000units/ml)-streptomycin (10mg/ml) 500ul, and B27 (1ml). Coverslips were washed over night in 1N nitric acid, washed 10x in sterile dH2O and stored in 200-proof ethanol until use. Prior to plating, coverslips were washed 3x in dH2O, and then coated overnight in 0.1mg/ml Poly-D-Lysine and 0.01mg/ml laminin in sterile dH20. For fixed analyses, plating medium was removed using a vacuum aspirator and neurons were washed 3x in warm HBSS. Neurons were then treated with 4% PFA in PHEM solution for 10 mins at RT. PFA was removed and neurons were washed 3x in HBSS and underwent immunohistochemistry staining protocols.

#### Microscopy

An inverted LSM 700 confocal microscope was used for imaging. For fixed brain sections, a complete Z-Stack was taken such that a single neuron was completely captured in a 2×2 tiling grid at a resolution of 1024 x 1024 pixels using a 20x objective. Entire somatosensory cortices labeled via IUE were imaged. For fixed cortical cultures, Z-stacks were taken to encapsulate an entire neuron. Tiling varied based on length of single neurons imaged.

#### Neuron tracing using SNT_analyzer.py

For our *in vivo* candidate screen, we generated 250um serial vibratome sections encompassing the entire primary somatosensory cortex (S1) from each brain and time point of interest. This section thickness enables acquisition of an entire layer II/III neuron axon spanning all cortical layers. Labeled neurons were imaged from the S1 region using a Zeiss 700 inverted confocal microscope. A complete z-stack encompassing the entire neuron was acquired in a 2×2 grid using a 20x objective. Using the entire z-stack of images, individual neurons were then traced in FIJI using the <u>S</u>imple <u>N</u>eurite <u>T</u>racer plugin (SNT) (Longair et al., 2011). In order to be included in the analyses, neurons must satisfy specific criteria: 1. The axon must span layers II-IV and reach layer V; and 2. The neuronal soma must reside within layer II/III. To determine whether the neuronal soma resided within layer II/III, prior to performing neuronal process tracing, the boundaries of layer IV were demarcated using DAPI staining to set these boundaries (Hand et al., 2015; Kelly and Hawken, 2017). Previous work has shown that layer II/III neurons have different branching patterns based on their soma location relative to layer IV (Larsen and Callaway, 2006). Specifically, layer II/III neurons with somas close to layer IV branch within layer IV, whereas those with somas further from layer IV branch in layer IV less often. Therefore, to ensure we included only the population of layer II/III neurons that rarely branches in layer IV (especially the middle region of layer IV), we plotted the soma distance from the border to layer IV versus the number of axon branches observed per neuron. We then used a linear regression model to determine a cutoff soma distance to be representative of our neurons of interest. For this, we assumed that the layer II/III neurons of interest do not branch in the middle of layer IV, we then interpolated the soma distance cutoff from the linear regression analysis. For P14 neurons, we calculated a y-intercept of 50.817 and a 95% confidence interval of (39.0μm–62.6μm). A cut-off of 39μm was used such that neurons with cell bodies greater than 39μm from the boundary to layer IV were scored (see Fig 2*H*). Neurons with cell body positions less than 39μm from the layer IV border were excluded from the data set with the assumption that their identity is ambiguous as to whether they are layer II/III neurons that normally branch in layer IV, or even perhaps a layer IV neuron. This ambiguity in part likely results from the arbitrary demarcation of layer IV boundaries using DAPI staining. For P21 neurons, we calculated a y-intercept of 79.9μm and a 95% confidence interval of (70.8μm-88.9μm). A cut-off of 71μm was used such that neurons with cell bodies greater than mm from the boundary to layer IV were scored. Neurons that met the “soma distance from layer IV” cut off criteria were then completely traced using the FIJI ImageJ SNT plugin. CSV files for each image were generated and fed into a python script (SNT_Analyzer.py). For layer V analyses at P14, we did not trace axons, but counted total primary branches formed in LV only in images where the entire extent of the axon in LV was visible. For neurons at P21, a significant number of traced axons were cut off in layer V during sectioning due to the larger axial spread at this later developmental time point. However, we observed that neurons whose primary axons could be traced for less than 125μm in layer V had a greater than 90% likelihood of displaying 0 primary branches in layer V, which is likely not an accurate picture of total branching in layer V. Therefore, we included only those neurons with primary axons that were traceable over a length longer than the 125μm cutoff in layer V. For cultured neurons, CSV files were generated and fed into a python script (SNT_Culture_analyzer.py). These programs enable computational analyses of neuron traces and ensures reproducibility by eliminating human error inherent in computing by hand. This code can be easily modified and or augmented to adjust for other calculations of interest, providing a valuable tool for the field.

#### Experimental design and statistical analyses

All graphs Prism version 9 (GraphPad). Student’s t-tests and analyses of variance (ANOVAs) comparisons were performed in Prism. The threshold for statistical significance was defined as p < 0.05. Data are presented as mean ± SEM, with n indicating the number of mice or neurons used in each group for comparison and is outlined below in Results. For *in vivo* OE and LOF IUE experiments (Figs. 2-5,7-8), an unpaired student’s t-test was used. For Dbn1 rescue experiments, a two-way ANOVA using Tukey’s multiple comparisons tests was used (see Fig. 9 for more details), Soma cutoff determination used for *in vivo* analysis is explained in Fig. 1. For *in vitro* OE and LOF experiments, an unpaired student’s t-test was used.

**Figure 1.**
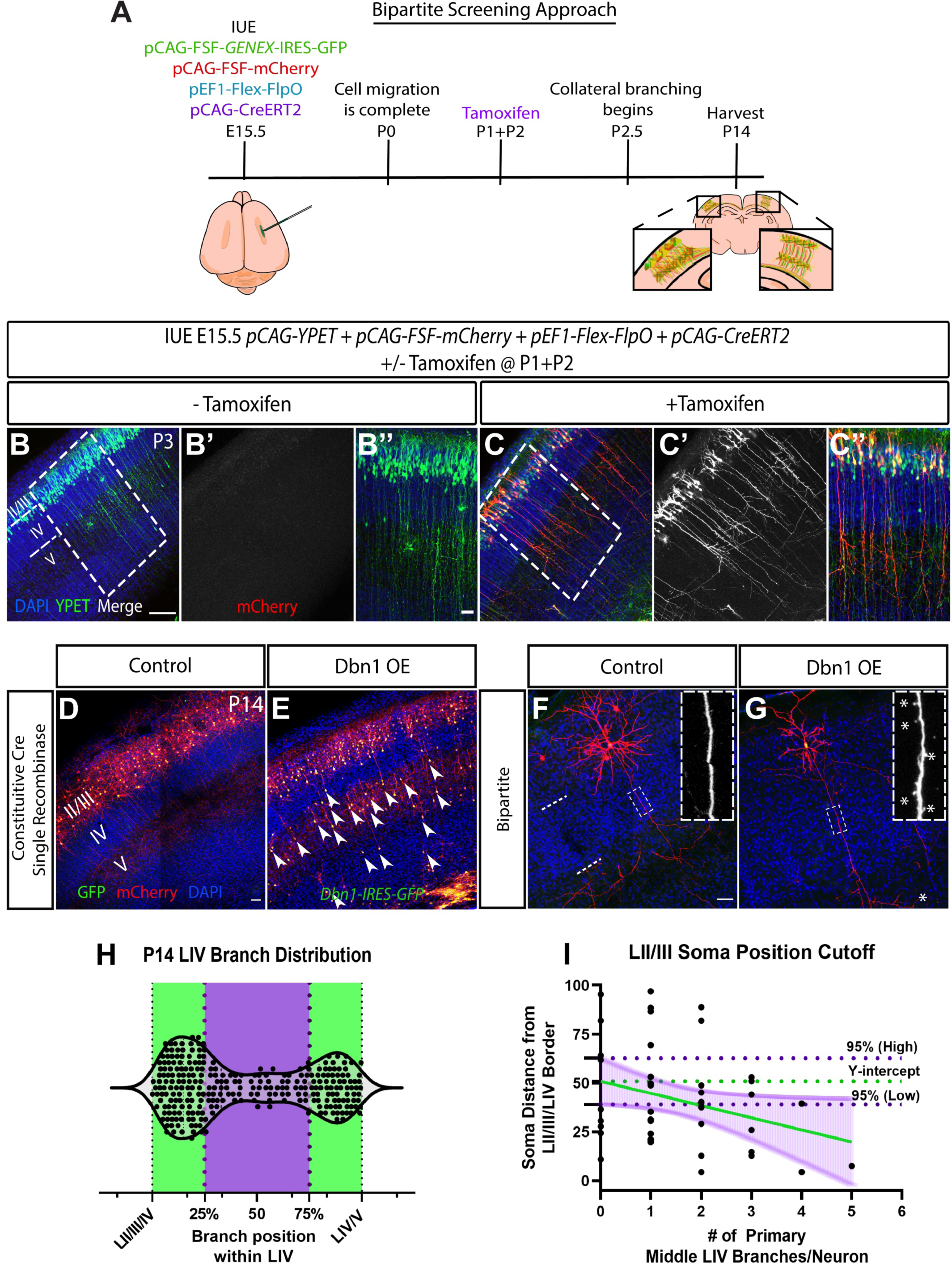
The bipartite *in utero* electroporation approach enables sparse and robust labeling of layer II/III cortical neurons *in vivo*. ***A,*** Schematic of the “Bipartite Screening Approach.” Inducible Cre recombinase and plasmids encoding Frt-Stop-Frt cassettes are used. ***B-C’’,***WT layer II/III neurons electroporated with *pCAG-YPET* (1ug/ul), *pCAG-FSF-mCherry* (1ug/ul), *pEF1-Flex-FlpO* (500ng/ul), and *pCAG-CreERT2* (500ng/ul). ***B-B’’,*** Confocal fluorescent maximum intensity projection images of WT CD1 layer II/III neurons from P3 pups that did not receive tamoxifen. Scale bar represents 100μm. ***B,*** Layer II/III neurons are labeled with *pCAG-YPET*. ***B’,*** No mCherry signal is detected since Cre is not recombined without addition of tamoxifen. ***B’’,*** Close up of cortical region outlined in ***B*** by dashed box. Scale box represents 25μm. ***C-C’’,*** Confocal fluorescent maximum intensity projection images of layer II/III neurons from P3 pups that received tamoxifen at P1 and P2. ***C,*** Layer II/III neurons are labeled with YPET and mCherry. ***C’,*** mCherry signal is present in layer II/III neurons. ***C’’,*** Close up of cortical section demarcated in ***C*** by dashed box. ***D-G,*** Example confocal maximum intensity images of somatosensory coronal section from P14 brains from the two screening approaches. ***D-E***, Examples using the constitutive Cre recombinase. Layer II/III neurons were electroporated with *pCAG-LSL-mCherry* (1ug/ul) (cell fill), *pCAG-LSL-GFP* (2ug/ul) (D), *pCAG-LSL-Dbn1-IRES-GFP* (2ug/ul) (E), and *pCAG-Cre* (500ng/ul) at E15.5. DAPI staining was used to distinguish cortical layer boundaries. ***D*,** Control neurons show typical laminar patterning. Scale bar represents 100μm. ***E,*** Dbn1 over-expression (OE) neurons show radial migration deficits (arrows). ***F-G,*** Examples using the “Bipartite” approach. Layer II/III neurons were electroporated with *pCAG-FSF-mCherry* (1ug/ul), *pCAG-FSF-GFP (*2ug/ul) (E), *pCAG-FSF-Dbn1-IRES-GFP* (2ug/ul) (F), and *pCAG-CreERT2* (500ng/ul) at E15.5. Tamoxifen was administered at P1 and P2. ***F,*** Single control neuron showing typical layer II/III pyramidal neuron branching morphology. Dashed lines represent cortical layer boundaries. Scale bar represents 100μm. Inset scale bar represents 5μm. ***G,*** Single Dbn1 OE neuron. No radial migration phenotype is observed. Boxed region highlights area of ectopic protrusions (asterisk). ***H,*** Volcano plot depicting the distribution and density of axon branches within layer IV as assessed in P14 cortices. Each dot represents an individual branch from a layer II/III neuron. Data were collected from 279 branches observed in 50 neurons across 3 independent electroporation experiments. Branch position percentage was calculated by measuring the entire length of the axon and determining where the branch fell with respect to soma position. Green areas denote branch positions in the superficial and deeper quarters of layer IV. ***I,*** Graph showing the distribution of layer IV primary branches versus soma distance from layer II/III/IV boundary. Each black dot represents a single neuron. Data were captured from 50 neurons across 3 independent electroporation experiments. We then used a linear regression model to determine a soma distance cutoff from the Layer II/III boundary for the purposes of scoring layer II/III neurons for branching parameters. We used the assumption that layer II/III neurons do not branch in the middle of layer IV. Therefore, for this linear regression, the Y-intercept, where Xi = 0 branches, is 50.817. The 95% confidence interval for the intercept is (39.00866–62.6247634). Therefore, we used the cutoff of layer II/III neurons with somas greater than 39μm from the layer II/III/layer IV border for scoring P14 neurons. The same calculations were performed for P21 neurons. The Y intercept where Xi = 0 branches is approximately 79.9μm with a 95% confidence interval range of (71-88). We therefore analyzed neurons with somas greater than 71μm from the layer II/III/layer IV border for scoring P21 neurons.

#### Code Accessibility

Python scripts for SNT-analyzer are publicly accessible via github: https://github.com/Jdors1/SNT-Analyzer.

## Results

### Visualizing individual cortical excitatory neuron morphology using *in utero* electroporation sparse labeling

Layer II/III excitatory pyramidal CPNs extend interstitial collateral axon branches mainly in layers II/III and V, both ipsilaterally and contralaterally (Greig et al., 2013; Hand et al., 2015). To target the layer II/III neurons that populate the somatosensory cortex, we performed *in utero* electroporation (IUE) at embryonic day 15.5 (E15.5). We employed a Cre recombinase-dependent mCherry construct that contains a STOP cassette flanked by *LoxP* sites (*Lox-Stop-Lox*, “*LSL*”) to label cell bodies and neurites and assess branching phenotypes at postnatal day 14 (P14), when cortical neuron process lamination is well established and potential interstitial axon branching phenotypes can be assessed (Hand et al., 2015). This enables sparse labeling by titrating the concentration of a co-electroporated Cre expression plasmid. We chose axon guidance receptors, signaling molecules and cytoskeletal regulators known to play roles in axon branching elsewhere in the nervous system and designed dominant-negative (DN), over-expression (OE) DNA constructs, shRNAs and guide RNAs (gRNAs) to perturb their function in and performed an *in vivo* IUE candidate screen.

In our initial experiments, using this “Single Recombinase Approach” described above, constructs were active from the time of electroporation (E15.5) and produced cortical neuron migration defects. One example is the migration defects observed after OE of the cytoskeletal regulatory protein drebrin (Fig. 1*D,E*). Since the regulation of cytoskeletal dynamics underlies both cell migration and axon branching, we hypothesized that constitutive Cre-driven gene-perturbations could preclude an assessment of laminar-specific interstitial axon branching (Cooper, 2013; Kapitein and Hoogenraad, 2015). Therefore, we bypassed these migration phenotypes by using a tamoxifen-inducible Cre construct (*pCAG-CreERT2)*. Since layer II/III CPN radial migration is complete by P0 and axon collateral branch initiation begins at P2.5 (Franco et al., 2011; Hand et al., 2015; Yoshinaga et al., 2021), we administered tamoxifen at P1 and then again at P2 using CreERT2. Use of CreERT2 in this experimental paradigm overcame cortical neuron migration phenotypes resulting from the use of constitutively active Cre in combination with various DNA constructs. However, we were unable to obtain robust neuronal labeling and had difficulty efficiently controlling the levels of Cre-mediated recombination (*data not shown*). We hypothesized that when using CreERT2, the low levels of Cre-mediated recombination and subsequent GFP expression were due to the short half-life of tamoxifen (Reid et al., 2014; Valny et al., 2016).

To overcome these issues, we developed a “Bipartite Screening Approach” (BP) (Fig. 1*A*) that uses inducible Cre-dependent DNA expression constructs that are not reliant on continuous tamoxifen administration. We employed a combination of Cre recombinase-dependent and FlpO recombinase-dependent DNA expression constructs (Fig.1*A*). We designed plasmids containing genes of interest embedded in a *Frt-STOP-Frt (FSF)* cassette instead of a *lox-STOP-lox (LSL)* cassette. We then delivered a combination of 3 or 4 plasmids (depending on the experimental paradigm) via IUE to layer II/III CPNs at E15.5 that included: *pCAG-FSF-mCherry* (cell fill), *pCAG-FSF-*GENEX*-IRES-GFP* (gene of interest), *pCAG-CreERT2*, and *pEF1-Flex-FlpO* (Fig. 1*A*). We also engineered an IRES-GFP into our gene expression constructs in order to monitor construct expression in electroporated cells. In this experimental paradigm FlpO expression is CreERT2-dependent. We administered tamoxifen at P1 and then again at P2, activating *pCAG-CreERT2* which in turn recombines *pEF1-Flex-FlpO*. FlpO is then constitutively “on” and can recombine the FlpO-dependent constructs to a much greater extent than we observed with a single tamoxifen-dependent Cre recombinase (Fig. 1*A* and *see below*). This approach allows for temporal control by employing a tamoxifen-dependent Cre recombinase, but also produces robust and sparse expression by titrating the expression of a Cre-dependent FlpO recombinase.

### Robust and sparse labeling of layer II/III neurons and visualization of collateral branching *in vivo*

Our Bipartite Screening Approach (BP) enables robust and sparse labeling of layer II/III neurons post-migration and satisfies the challenge of obtaining single cell resolution of cortical neuron axons and their branches to assess collateral branching phenotypes *in vivo*. To validate our method, we first performed bulk labeling of layer II/III neurons using a *pCAG-YPET* DNA construct in addition to the other BP constructs, but without tamoxifen treatment. YPET is a Yellow Fluorescent Protein (YFP) modified for FRET imaging, which we have found is robustly detected using antibodies directed against GFP in our electroporation paradigm and thus was used for these initial experiments (Nguyen and Daugherty, 2005) (labeled as “GFP” in Fig. 1 panels). By P3, just after the onset of axon branch initiation, YPET-labeled layer II/III neurons identified by immunohistochemistry (IHC) have cell bodies that populate the upper layers of the cortex (layers II and III) and begin to extend axon branches in layer V (Fig. 1*B-B’’*). However, we could not visualize individual axon branches in detail using bulk labeling in conjunction with the BP approach without tamoxifen (Fig. 1*B’’*). Importantly, IUE with this combination of plasmids in the absence of tamoxifen is not leaky, since IUE of FlpO-dependent mCherry in combination with the other constructs does not result in mCherry expression (Fig. 1*B’*). However, tamoxifen administration in conjunction with IUE of BP plasmids results in sparse and robust mCherry labeling of axons, allowing for detailed assessment of collateral branches across all cortical layers at the single cell level (Fig. 1*C–C’’*). Together, these results demonstrate that the BP approach allows for: (i) temporal regulation of labeling and perturbation constructs; (ii) segregation of neuronal migration from axon branching phenotypes; and (iii) sparse layer II/III neuron labeling with the ability to titrate the number of neurons labeled without sacrificing robust gene expression.

Using the BP approach, we analyzed the distribution of cortical layer II/III neuron interstitial axon branches across all cortical layers in wild type mice at P14. We found that layer II/III CPNs do show a modest degree of collateral axon branching in layer IV, however, these branches mainly occur at the boundaries between layers II/III and IV (Fig. 1*H-I*). We segmented layer IV in order to determine the distribution of branching within this layer, and we observed that very little axon branching occurs in the middle half of layer IV (Fig. *1H*). Interestingly, we observe a correlation between cell-body depth in layer II/III with the number of axon collaterals formed in middle layer IV; neurons closer to the layer II/II-IV boundary exhibit collateral branches at a greater frequency within the middle of layer IV than II/III neurons with cell bodies closer to layer I (Fig. 1*I*). These observations are in line with previous findings showing that layer II/III CPNs with cell bodies located more superficially lack axon arbors in layer IV, while CPNs whose cell bodies are closer to the layer IV border exhibit substantial axon branching within layer IV in the developing mouse brain (Larsen and Callaway, 2006). Together, these results highlight the existence of distinct classes of layer II/III neurons and support the idea that soma position of these neurons predicts axonal branching patterns.

Using these data we identified a cutoff soma position to exclude layer II/III neurons that reside close to the border of layer IV and so are more prone to forming collateral branches in layer IV (Fig 1*I*; see Methods). We chose to focus here only on layer II/III neurons that normally do not branch in layer IV in our search for determinants of laminar specific interstitial axon branching patterns.

### Dbn1 over-expression leads to an increase in axon protrusions in layer II/III CPNs

We initially screened candidate genes encoding cytoskeletal regulators using our constitutive Cre-dependent Single Recombinase Approach. Among these candidates were Dbn1, the Ras-family GTPase RalA, and the serine/threonine kinase GSK3β (Lalli and Hall, 2005; Kim et al., 2011; Ketschek et al., 2016). Over-expression of WT Dbn1 using the Single Recombinase Approach resulted in significant layer II/III radial neuronal migration deficits as assessed at P14 (Fig. 1*E*). Because over-expression of several candidate genes, including Dbn1 and also a constitutively active-RalA (CA-RalA: RALAG23V (Lopez et al, 2008)), resulted in severe layer 2/3 cortical neuron migration defects, we re-screened these candidates using the BP approach (Fig. 1*F-G*, 2*A-D’’*).

Using the BP approach, we found that both CA-RalA and CA-GSK3β showed the presence of ectopic axon branches with aberrant trajectories in layer IV, demonstrating that bypassing CPN migration defects and manipulating genes of interest just prior to the onset of collateral branching enables identification of branching defects (Fig. 2*C-C’’,D-D’’,G,H*). Dbn1 OE, however, did not result in an increase in total axon collateral branches within layer IV, but instead led to the formation of ectopic protrusions along the entire length of the axon (Figs. 2*B-B’’,F*). We define neurites or axon collaterals less than 10 micrometers (μm) in length extending from the main CPN axon shaft as protrusions (Ketschek et al., 2016). These protrusions resemble our previous observations showing in WT layer II/III CPNs at P3 that axon collateral initiation events are uniformly distributed across all cortical layers; however, later in the development of layer II/III CPNs, a subset of protrusions in layers II/III and V extend to form long and stable collaterals, while those in layer IV are retracted (Hand et al., 2015). Additional studies also show that filopodial axon protrusions serve as the precursors for axon branches (Gibson and Ma, 2011; Hand et al., 2015; Ketschek et al., 2016; Armijo-Weingart and Gallo, 2017). Therefore, in Dbn1 OE CPNs, persistent axon protrusions at P14 provide evidence of collateral branch initiation events that fail to undergo later elongation or retraction.

**Figure 2.**
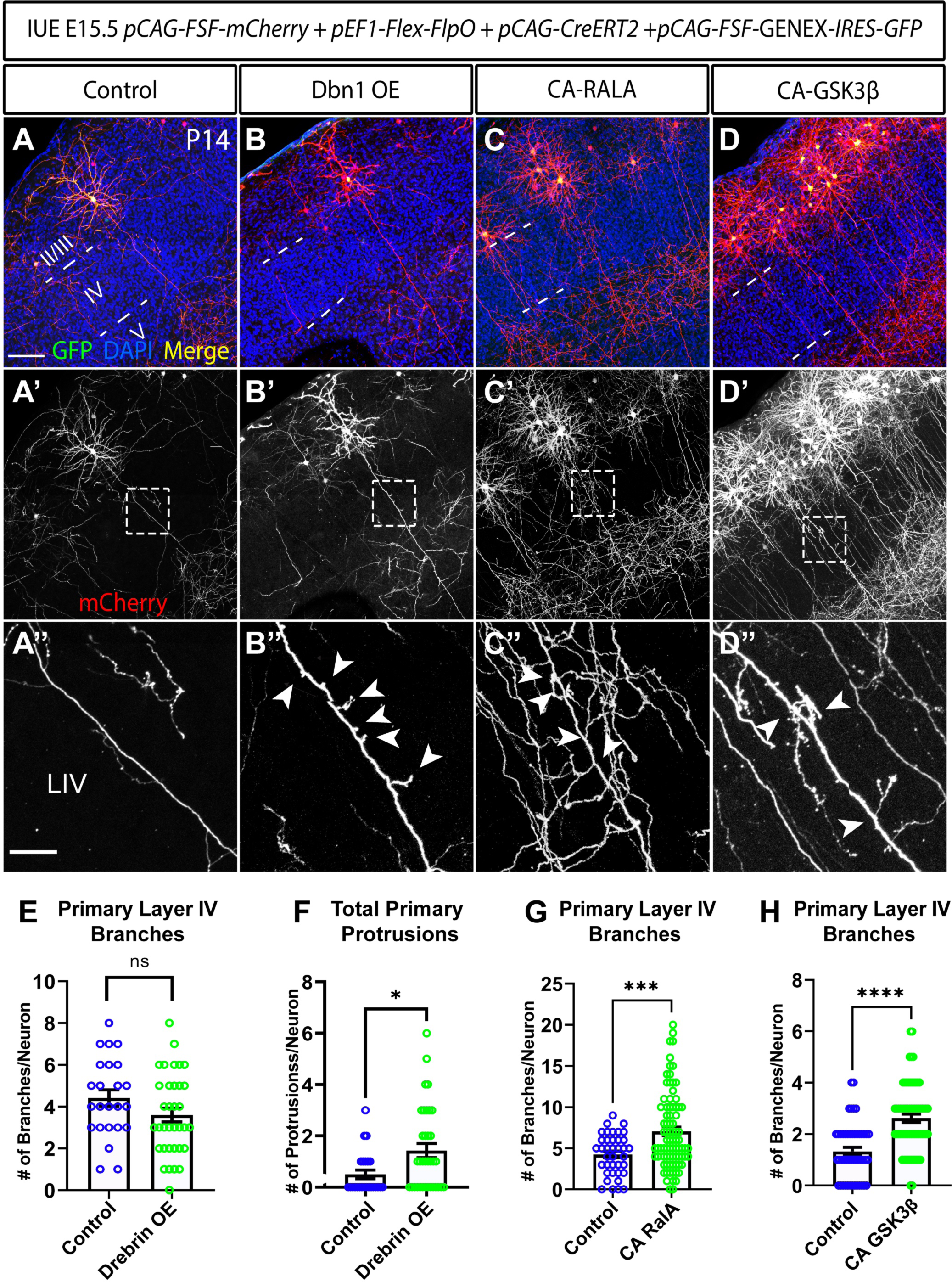
*Dbn1* over-expression in layer II/III neurons *in vivo* causes ectopic and stable protrusions along primary axons. ***A-D’’,*** Confocal immunofluorescent maximum intensity projection images of layer II/III P14 neurons from CD1 WT mice electroporated with over-expression candidates from the Single Recombinase/Constitutive Cre approach using the bipartite labeling approach. All neurons were electroporated with: *pCAG-FSF-GENEX-IRES-GFP* (2ug/ul), *pCAG-FSF-mCherry* (1ug/ul), *pEF1-Flex-FlpO*, and *pCAG-CreERT2*. Pups were administered tamoxifen at P1 and P2 (1mg/ml). Brains were harvested and sectioned at P14 and immunostained with antibodies directed against GFP, against dsRED to detect mCherry, and DAPI to assess potential branching patterns that may have been masked by migration deficits in the original screening approach (these same staining conditions are used throughout all following figures except where specified). ***A,*** Control neurons were electroporated with *pCAG-FSF-GFP* (2ug/ul) in addition to other 3 bipartite constructs. Scale bar represents 100μm. Littermate controls were used for statistical calculations within each experiment ***A’,*** mCherry labeling of electroporated neurons (cell fill). ***A’’,*** Zoomed in view of middle layer IV axon region. Scale bar represents 25μm. ***B,*** *Dbn1* OE. Neurons were electroporated with *pCAG-FSF-Dbn1-IRES-GFP* in addition to other 3 bipartite constructs. ***B’,*** mCherry labeling of electroporated neurons. ***B’’*,** Zoomed in view of layer IV region of layer II/III axon. ***C-C’’,*** CA-RALA. ***C,*** Neurons were electroporated with *pCAG-FSF-CA-RALA-IRES-GFP* in addition to other 3 bipartite constructs. ***C’,*** mCherry labeling of layer II/III neuron. ***C’’,*** Zoomed in view of layer IV region along layer II/III neuron axon. ***D-D’’,*** CA-GSK3β. ***D,*** Layer II/III neurons were electroporated with *pCAG-FSF-CA-GSK3β-IRES-GFP* in addition to other 3 bipartite constructs. ***D’,*** mCherry labeling of layer II/III neurons. ***D’’,*** Zoomed in view of layer IV region along layer II/III axon. ***E-I,*** Quantification of protrusion and branching phenotypes. ***E,*** Total number of branches within layer IV for *Dbn1* OE neurons. No significant difference was found in the number of branches between control and OE neurons. ***F,*** Total number of protrusions for *Dbn1* OE layer II/III neurons across all layers. Dbn1 OE causes an increase in the total number of axon protrusions (branches <10μm). ***G,*** Total number of layer IV branches (>10μm). Over-expression of CA-RalA causes an increase in the number of branches in layer IV. 2 independent electroporations with 2-3 pups per conditions were analyzed. N=38 control neurons and 83 CA-RALA neurons. ***H,*** Over-expression of CA-GSK3β causes an increase in axon branches in layer IV. 3 independent electroporations with 2-3 pups per condition per experiment were analyzed. N=64 control neurons and N=66 CA-GSK3β neurons. Statistics: unpaired t-test. Lines represent mean and SEM. Symbols represent individual neurons obtained from independent experiments. *p < 0.05, **p<0.01, ****p<0.0001.

Septin7, like Dbn1, plays a role in regulating the axon cytoskeleton during neurite outgrowth (Hu et al., 2012; Kaplan et al., 2017; Radler et al., 2023). Septin7 binds to MTs and promotes the entry of axonal MTs into filipodia during the initial steps of collateral branching, similar to what is thought to be the mechanism underlying Dbn1 function (Hu et al., 2012; Ketschek et al., 2016). We assessed *Septin7* overexpression using the BP approach, but we did not observe any changes in axon branching or protrusions in layer II/III CPNs (*data not shown*). This suggests that *Dbn1* overexpression phenotypes are specific to drebrin and not a general property of this class of cytoskeletal binding proteins.

Taken together, these data show that *Dbn1* overexpression causes an increase in stable axon protrusions in layer II/III CPNs throughout cortical development. *Dbn1* overexpression did not result in a change in the number of collateral axon branches extending from the primary axon, suggesting that these aberrant protrusions do not develop into stabilized, mature, collateral branches.

### Dbn1 loss-of-function in layer II/III neurons results in ectopic collateral axon branching within layer IV

We next analyzed how LOF in genes encoding cytoskeletal regulatory proteins of interest in this study affects the formation of stable axon collateral branches. For *RalA* LOF, we over-expressed a constitutively inactive GDP-bound RalA (G26A) (Vitale et al., 2005) in layer II/III neurons using the BP approach and did not see any abnormal branching phenotypes (*data not shown)*. We also used the BP approach to over-express a dominant negative GSK3β (K85A) (Dominguez et al., 1995) and did not observe any branching phenotypes (*data not shown)*. However, based on the strong collateral axon branching phenotypes we observed following *Dbn1* LOF in layer II/III CPNs (see below), we focused on drebrin.

To assess *Dbn1* LOF in layer II/III CPNs, we used the BP approach in the context of a *Dbn1^flox^* conditional allele (Zhang et al., 2018), leading to *Dbn1* LOF at ∼P3, the normal onset of collateral axon branch initiation. Since *Dbn1* OE results in an increase in collateral branch initiation (manifested as protrusions) but not collateral axon branch stabilization (mature axon branches), we hypothesized that *Dbn1* LOF would result in a decrease in protrusions and an overall reduction in layer II/III neuron axon branches. In heterozygous *Dbn1^fl/+^* littermate control mice subjected to BP IUE there were very few to no collateral axon branches in layer IV, including within the middle region of layer IV (Fig. 3*A-A’’*). However, *Dbn1* LOF, assessed using the BP approach in *Dbn1^fl/fl^*mice, resulted in an increase in collateral axon branching in layer II/III neurons within layer IV compared to heterozygous controls observed at P14 (Figs. 3*B-F*). This increase was seen within the middle region of layer IV, where control neurons rarely branch, and therefore the presence of increased numbers of collateral axon branches in this region, even just one or two, is statistically significant. There was no difference in collateral axon branching along the primary axon within layer II/III or layer V following *Dbn1* LOF in layer II/III CPNs (Fig. 3*H,I*). In contrast to *Dbn1* OE in layer II/III neurons, *Dbn1* LOF did not produce differences in the number of primary axon protrusions within layer IV (Fig. 3*G*). Therefore, *Dbn1* OE and LOF in layer II/III neurons lead to distinct phenotypes; overexpressing Dbn1 causes an increase in axon protrusions but not stabilized axon collateral branches, whereas *Dbn1* LOF in layer II/III neurons results in ectopic axon collateral branches in layer IV.

**Figure 3.**
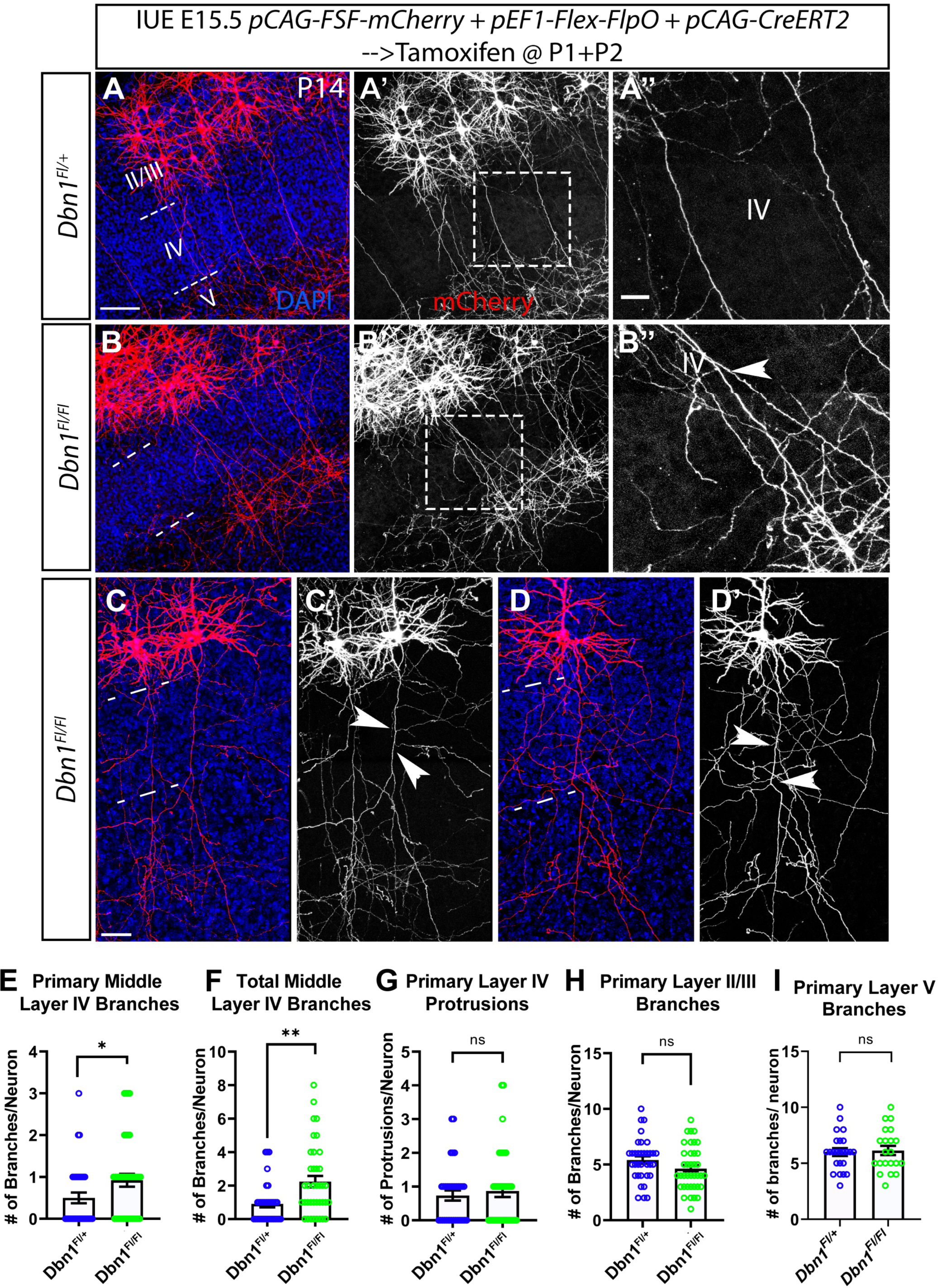
*Dbn1* LOF in layer II/III neurons *in vivo* results in ectopic branching in the middle of layer IV. ***A-D’,*** Confocal immunofluorescent maximum intensity projection images of layer II/III P14 neurons from *Dbn1* floxed and heterozygous littermate controls on a mixed background labeled via electroporation with *pCAG-FSF-mCherry* (1ug/ul), *pEF1-Flex-FlpO* (500ng/ul), and *pCAG-CreERT2* (500ng/ul) using the bipartite approach. Tamoxifen was administered at P1 and P2. ***A-A’’,*** Example of *Dbn1* heterozygous layer II/III LOF neurons. ***A,*** Merged image with scale bar that represents 100μm. ***A’,*** mCherry labeling. ***A’’,*** Zoomed in example of layer IV region along layer II/III neuron axon. Scale bar represents 50μm. ***B-B’’,*** Example of *Dbn1* mutant layer II/III neuron. ***B’,*** mCherry labeling of mutant neuron. ***B’’,*** Zoomed in the middle layer IV region along layer II/III axon. Arrows point to the presence of ectopic branches in this region. ***C-D’,*** Examples of neurons that show dramatic ectopic branching in layer IV due to loss of Dbn1. ***C’,D’,*** mCherry staining of neurons. Arrows point to ectopic branches in the middle layer IV region. Dashed lines demarcate layer borders. ***E-I,*** Quantification of layer II/III neuron axon branches and protrusions. Symbols represent individual neurons obtained from: Fl/+: 34 individual neurons from 5 mice obtained from 2 independent experiments, Fl/Fl: 38 individual neurons from 4 mice obtained from 2 independent experiments. For layer V quantification, 24 Fl/+ and 21 Fl/Fl neurons were analyzed. Statistics: unpaired t-test. Lines represent mean and SEM. *p<0.05, **p<0.01, ***p<0.001, ****p<0.0001.

To verify the requirement for Dbn1 in suppressing ectopic axon collateral branching, we performed rescue experiments using the *Dbn1^flox^* allele to provide cell-autonomous loss of Dbn1 in layer II/III CPNs. We introduced plasmids to express *Dbn1* in *Dbn1^fl/+^* and *Dbn1^fl/fl^* layer II/III CPNs using a slightly modified BP approach (see Methods) and assessed axon branching phenotypes at P14. We found, again, that *Dbn1* LOF layer II/III neurons resulted in ectopic branches in the middle region of layer IV with no presence of ectopic protrusions in this region (Fig. 4*A-B’’*). *Dbn1* OE in *Dbn1^fl/fl^* layer II/III neurons rescued the *Dbn1* ectopic branching LOF phenotype in the middle region of layer IV, (Fig. 4*B-B’’,G*). *Dbn1* expression in *Dbn1^fl/+^* but not in *Dbn1^fl/fl^* layer II/III CPNs (which lack Db11), resulted in an increase in axon protrusions within layer IV (Fig. 4*B-B’’*,*F*), further indicating that high levels of Dbn1 levels result in protrusions. Using a construct expressing YFP-tagged Dbn1, we also observed localization of Dbn1 within CPN axons in layer IV (Fig. 4*B’’*). These data support the conclusion that Dbn1 suppresses the formation of collateral axon branches in the middle region of layer IV.

**Figure 4.**
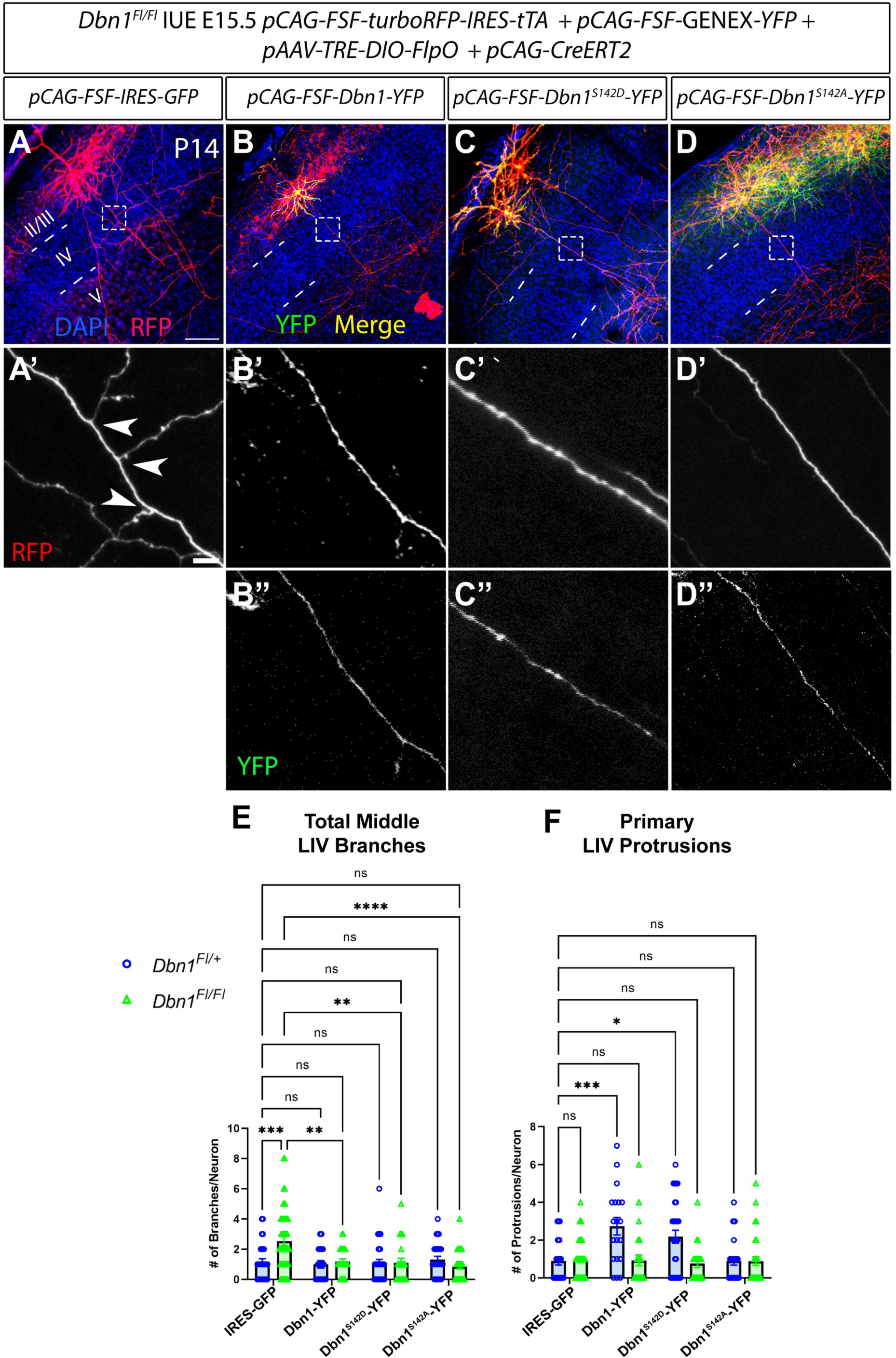
Phosphorylation of Dbn1 at S142 is not required for Dbn1 suppression of ectopic collateral branching in layer IV. ***A-D’’*,** Confocal immunofluorescent images of layer II/III neurons at P14 obtained from *Dbn1^flox^* mice using *ex vivo* electroporation at E15.5. Mice were electroporated using a modified version of the bipartite approach with the following constructs: *pCAG-CreERT2* (0.3-0.5μg/ul), *pAAV-TRE-DIO-FlpO* (0.01-0.015μg/ul), and *pCAG-FSF-turboRFP-IRES-tTA* (0.75-1μg/ul). All (‘) letters represent zoomed in images of dashed box region in respective letter image with tagRFP staining. All (‘’) represent zoomed in images of dashed box region in respective letter image with GFP staining (for Dbn1 localization). ***A***, Layer II/III neurons were additionally electroporated with *pCAG-FSF-IRES-GFP* (1.5μg/ul). Scale bar represents 100μm. ***A’***, Scale bar represents 10μm. ***B-D’’,*** Layer II/III neurons were electroporated with an additional *Dbn1* expression construct: ***B-B’’***, *pCAG-FSF-Dbn1-YFP* (1.5μg/ul), ***C-C’’***, *pCAG-FSF-Dbn1^S142D^-YFP* (1.5μg/ul), ***D-D’’,*** *pCAG-FSF-Dbn1^S142A^-YFP* (1.5μg/ul). ***E-F***, Quantification of axon branches and protrusions for experimental perturbations. The following number of neurons were analyzed per condition: Fl/+: IRES-GFP (control): 28 neurons across 6 mice from 4 independent IUEs, Dbn1 WT: 19 neurons across 2 mice from 1 IUE, Dbn1^S142D^: 33 neurons across 2 mice from 1 IUE, Dbn1^S142A^: 28 neurons across 2 mice, from 2 independent IUEs. Fl/Fl: IRES-GFP: 50 neurons, from 6 mice from 2 independent IUEs, Dbn1 WT: 27 neurons across 5 mice from 2 independent IUEs, Dbn1^S142D^: 21 neurons across 4 mice from 2 independent IUEs, Dbn1^S142A^: 35 neurons across 3 mice from 2 independent IUEs. Statistics: Two-way ANOVA using Tukey’s multiple comparisons test. *p<0.01, **p<0.001, ***p<0.0001, ****p<0.00001.

### Ectopic collateral axon branching following *Dbn1* loss-of-function persists at later developmental stages

Since synaptic pruning and axon elimination can occur both early and later in postnatal development (Luo and O’Leary, 2005; Riccomagno and Kolodkin, 2015), we asked whether *Dbn1* LOF phenotypes are observed at more advanced postnatal developmental stages. We assessed axon branching in *Dbn1* LOF layer II/III CPNs at P21 *in vivo*. P21 *Dbn1^fl/fl^* layer II/III neurons do show ectopic branches in the middle region of layer IV that are typically absent in *Dbn1^fl/+^*layer II/III neurons (Fig. 5*A-E*). Further, *Dbn1* LOF CPNs do not show the presence of excess protrusions in layer IV (Fig. 5*F*). These data suggest that ectopic axon branches observed at P14 due to *Dbn1* LOF persist into late postnatal development. The absence of primary ectopic protrusions within the middle region of layer IV also suggests that no new ectopic branches are formed between P14 and P21. Similar to P14, *Dbn1* LOF CPNs exhibit no difference in the number of branches within layer II/III or V compared to *Dbn1^fl/+^* CPNs (Fig 5*G,H*). These data suggest Dbn1 restricts the growth of ectopic axon branches in the middle region of layer IV, where branching is normally absent, and that these effects persist and are not corrected.

**Figure 5.**
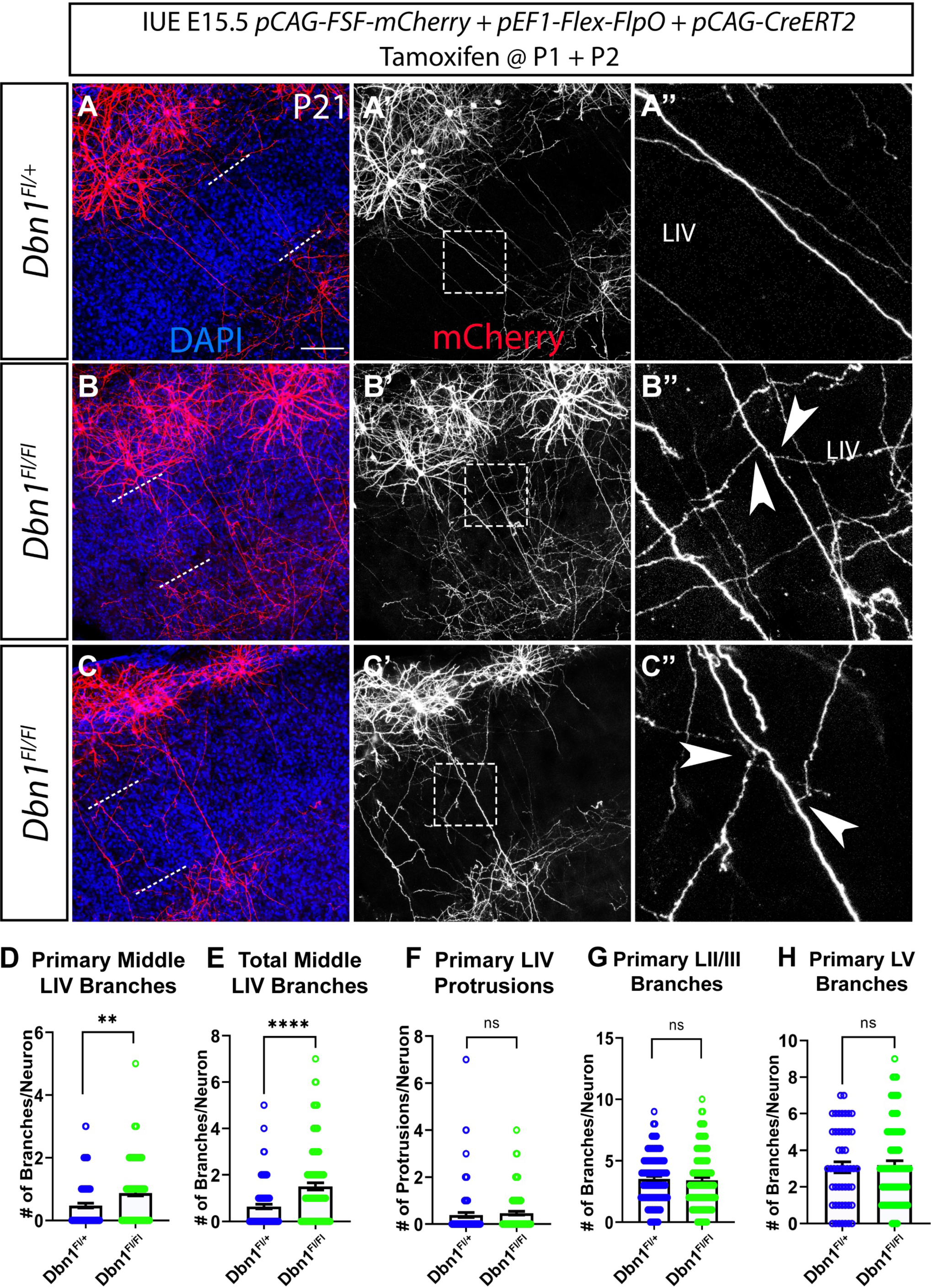
*Dbn1* LOF ectopic branching phenotypes are persistent throughout later postnatal developmental time points. ***A-B’’,*** Confocal immunofluorescent maximum intensity projection images of layer II/III P21 neurons from *Dbn1* floxed and heterozygous littermate controls labeled using the “Bipartite” approach with the following constructs: *pCAG-FSF-mCherry* (1ug/ul), *pEF1-Flex-FlpO* (500ng/ul), and *pCAG-CreERT2* (500ng/ul). Tamoxifen was administered at P1 and P2. ***A-A’’,*** Example heterozygous neuron. ***A,*** Merged image of mCherry and DAPI staining. Dashed lines demarcate cortical layer boundaries. ***A’,*** mCherry staining. Dashed box represent region along the axon within the middle of layer IV. ***A’’,*** Magnified image of boxed region in **A’**. Scale bar represents 25μm. ***B-B’’,*** Example of *Dbn1* null neuron. ***B,*** Merged image of mCherry and DAPI staining. Dashed lines demarcate cortical layer boundaries. ***B’,*** mCherry staining. Dashed box represents region along the axon within the middle of layer IV. ***B’’,*** Magnified image of boxed region in ***B’***. Arrows point to ectopic branches in layer IV. ***C-E,*** Quantification of layer IV branches and protrusions. ***F,*** Quantification of layer II/III branches. ***G,*** Quantification of layer V branches. ***H-I,*** Quantification of middle layer IV branches for pups receiving tamoxifen at later time points. Symbols represent individual neurons obtained from: Fl/+: 95 individual neurons from 4 mice obtained from 3 independent experiments, Fl/Fl: 110 individual neurons from 7 mice obtained from 3 independent experiments. For layer V analyses: 50 individual Fl/+ neurons and 85 individual Fl/Fl neurons obtained from the same experiments were used due to additional LV cutoff (see methods). Statistics: unpaired t-test. Lines represent mean and SEM. *p<0.05, **p <0.01, ***p<0.001, ****p<0.0001.

### The requirement for Dbn1 to regulate axon collateral branching is confined to an early postnatal developmental window

To determine whether ectopic collateral axon branching observed in Dbn1-deficient layer II/III CPNs is confined to the narrow developmental window around the time of collateral axon branch initiation (∼P2-P3) after cell migration is complete (P0), we performed LOF experiments in *Dbn1^flox^* mice, as described above, but administered tamoxifen at later postnatal stages (P4 and P7) (Fig. 6*A-D’*)*. Dbn1^fl/fl^* mice that received tamoxifen at later times, at either P4 or P7, and were subsequently sacrificed at P14 resembled *Dbn1^fl/+^* littermates, with layer II/III neurons exhibiting no ectopic collateral axon branching phenotypes in the middle region of layer IV (Fig. 5*E-F*). Therefore, *Dbn1* LOF CPN interstitial branching phenotypes are developmentally constrained such that only removal of Dbn1 just prior to the onset of collateral axon branching results in axon collateral branching phenotypes.

**Figure 6.**
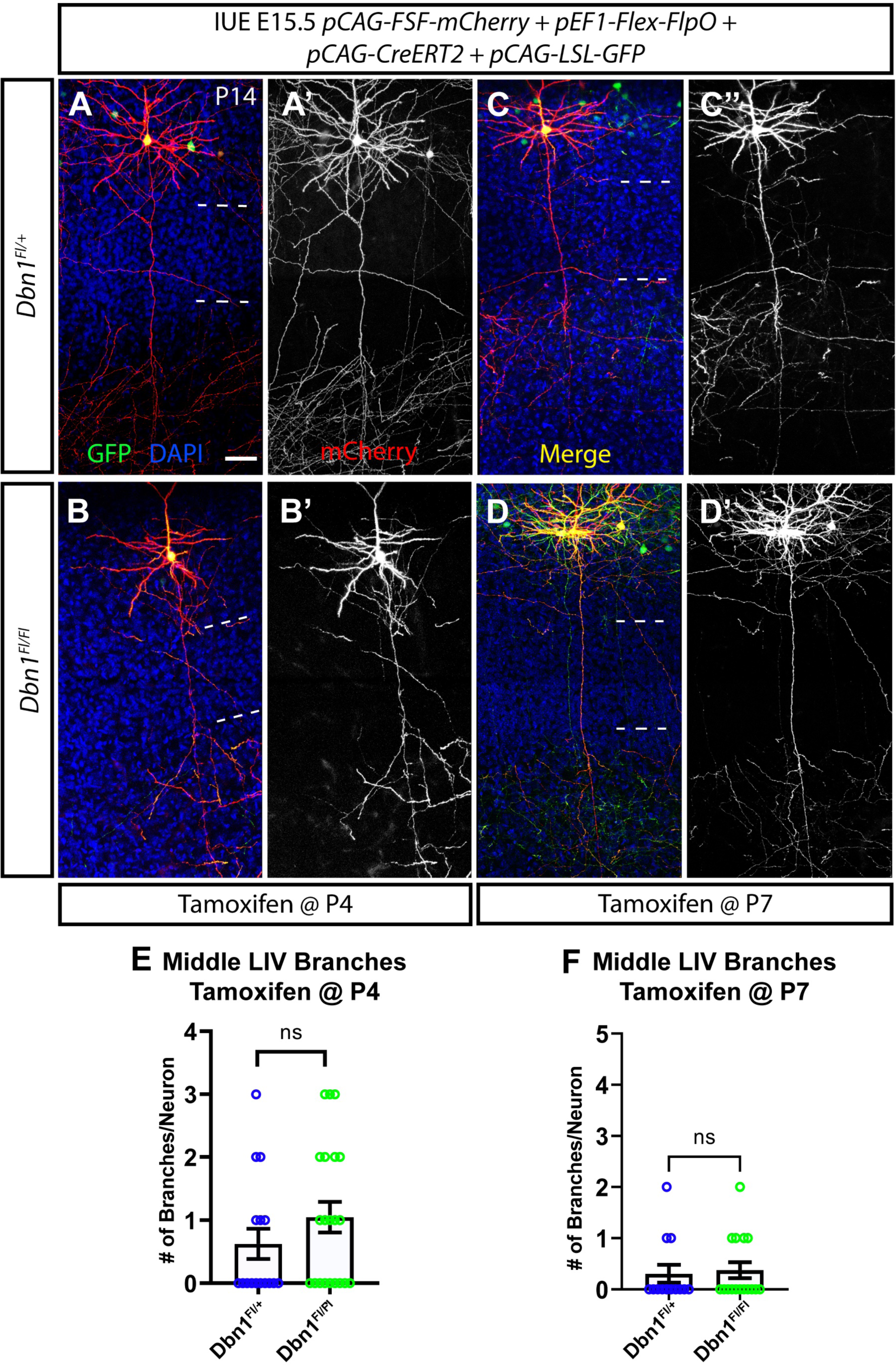
*Dbn1* LOF ectopic branching phenotypes are developmentally fixed. ***A-D’,*** Neurons were electroporated with *pCAG-LSL-GFP* (1ug/ul), *pCAG-FSF-mCherry* (1ug/ul), *pEF1-Flex-FlpO* (500ng/ul) and *pCAG-CreERT2* (500ng/ul). ***A-B’,*** *Dbn1^fl/+^* pups (A-A’) and *Dbn1^fl/fl^* pups (B-B’) were administered tamoxifen at P4 after branching has already initiated. Confocal fluorescent maximum intensity projection image of P14 cortical section stained with mCherry, GFP and DAPI. No branching defects were observed. Scale bar represents 50μm. ***C-D’,*** *Dbn1^fl/+^*pups (C-C’) and *Dbn1^fl/fl^* pups (D-D’) were administered tamoxifen at P7 several days after branching has already initiated. Confocal fluorescent maximum intensity projection image of P14 cortical section stained with mCherry, GFP and DAPI. No branching defects were observed. Dotted lines demarcate layer boundaries. ***E-F,*** Quantification of layer II/III neuron axon branches within the middle region of layer IV. For P4 tamoxifen treatment symbols represent individual neurons obtained from: 16 Fl/+ neurons from 2 mice and 21 Fl/Fl neurons from 2 mice obtained from 1 electroporation experiment. For P7 tamoxifen treatment symbols represent 13 Fl/+ neurons from 1 mouse and 16 Fl/Fl neurons from 2 mice obtained from 1 electroporation experiment. Statistics: unpaired t-test. Lines represent mean and SEM.

### Dbn1 over-expression and loss-of-function axon protrusion and branching phenotypes are recapitulated *in vitro*

Dbn1 over-expression in DRGs and in embryonic cortical neurons results in an increase in axon branches, filopodial protrusions and neurites *in vivo* and *in vitro* (Geraldo et al., 2008; Ketschek et al., 2016; Poobalasingam et al., 2022). Additionally, knock down of Dbn1 using RNAi leads to a reduction in filopodia, lamellipodia and axon branching *in vitro* in several neuronal cell types, including cortical neurons and sensory neurons (Geraldo et al., 2008; Ketschek et al., 2016; Miyata and Hayashi, 2022). To complement these findings and our *in vivo* results on interstitial axon branching regulators we manipulated layer II/III CNPs *in vitro*. We performed *ex vivo* electroporation on *Dbn1^fl/fl^* mouse embryos at E15.5 using plasmids expressing Cre recombinase (for *Dbn1* LOF) or only GFP (control) to label layer II/III neurons. Then, we immediately dissociated electroporated cortices, cultured isolated neurons, and subsequently assessed cortical neuron axon morphology after 6 days *in vitro* (DIV), a time point that corresponds to the postnatal stage at which we normally observe the onset of layer II/III collateral axon branching *in vivo* (∼P2; (Hand et al., 2015)). *Dbn1^fl/fl^* neurons electroporated with Cre exhibit a loss of Dbn1 protein compared to control neurons electroporated with GFP, demonstrating efficient inactivation of Dbn1 (Fig. 7*A-B’*).

**Figure 7.**
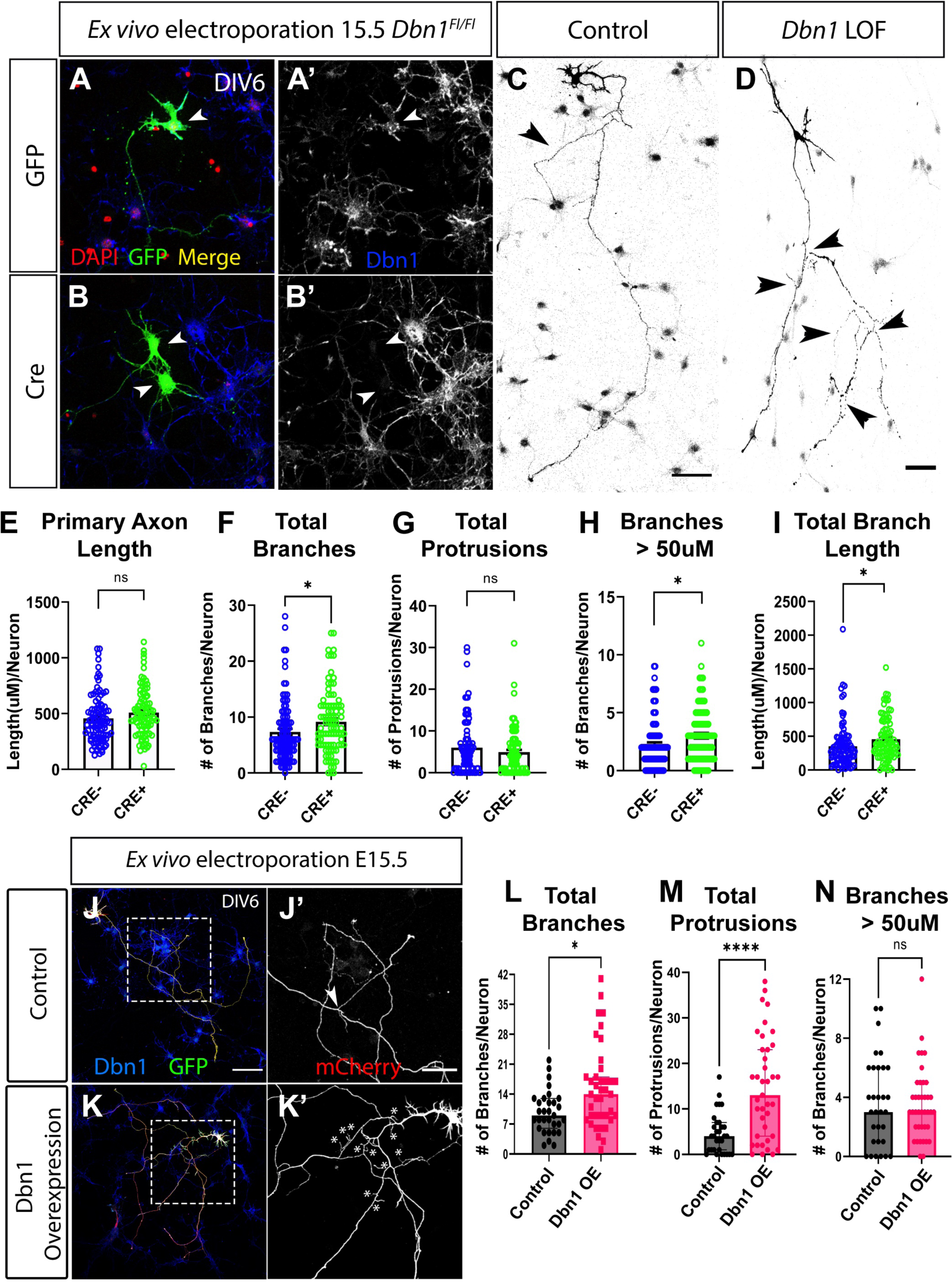
*Dbn1* LOF in layer II/III neurons *in vitro* results in longer and more axon branches. ***A-D,*** Confocal immunofluorescent images of layer II/III neurons at DIV6 obtained from *Dbn1^fl/fl^* mice using *ex vivo* electroporation at E15.5. Cells were stained with antibodies directed against GFP, against dsRED, against Dbn1, and DAPI for analyses. ***A,*** *Dbn1^fl/fl^* embryos were electroporated at E15.5 with *pCAG-GFP* (2ug/ul) (control neurons). Example of control neuron. Arrow points to GFP-positive soma. ***A’’,*** Dbn1 staining showing the presence of Dbn1 in the GFP labeled neurons. Arrow points to GFP-positive cell body. Boxed region highlights example of average branch size found in control neurons. ***B-B’,*** Neurons from *Dbn1^fl/fl^* were electroporated with *pCAG-LSL-GFP* (2ug/ul) and *pCAG-Cre* (1ug/ul) (LOF neurons). ***B,*** A GFP labeled *Dbn1* LOF layer II/III neuron soma. ***B’,*** Dbn1 staining shows the absence of Dbn1 protein in this GFP-positive neuron. Arrowhead indicates absence of Dbn1 in the GFP-positive cell body. Boxed region shows zoomed in image of average size of an axon branch found in LOF neurons. ***C-D,*** Example of Control (*Dbn1^fl/+^*) versus Dbn1 LOF (*Dbn1^fl/fl^*) neuron. Arrows point to axon branches. Scale bars represent 50μm. ***E-I,*** Quantification of axon branches. Control (-Cre): 95 neurons obtained from 2-3 pups across 2 individual experiments, LOF: (+Cre): 92 neurons obtained from 2-3 pups across 2 individual experiments. Statistics: unpaired t-test. Lines represent mean and SEM. *p<0.05, **p<0.01. ***J-K’,*** Confocal immunofluorescent images of layer II/III neurons at DIV6 obtained from WT CD1 mice using *ex vivo* electroporation at E15.5. Neurons were electroporated with *pCAG-FSF-mCherry* (1ug/ul), *pCAG-FSF-GFP* (J), *pCAG-FSF-Dbn1-IRES-GFP* (2ug/ul) (K), and *pCAG-FlpO* (1ug/ul). ***J,*** Example of WT control neuron. Dashed box represents region expanded in J’. Scale bar represents 100μm. ***J’,*** Expanded view of boxed region with dsRed staining. Scale bar represents 50μm. ***K,*** Confocal immunofluorescent images of Dbn1 OE layer II/III neurons at DIV6 obtained from WT CD1 mice using *ex vivo* electroporation at E15.5. Dashed box represents region expanded in K’. ***K’*** Zoomed in image of axon protrusions; asterisks indicate protrusions. ***L-N,*** Quantification of axon branches and protrusions. 46 control neurons and 41 Dbn1 over-expressing neurons were analyzed across 2-3 brains across 2 independent experiments (2 separate cultures). Statistics: unpaired t-test. Lines represent mean and SEM. *p<0.05, **p<0.01, ***p<0.001, ****p<0.0001.

*Dbn1* LOF in cultured layer II/III neurons resulted in an increase in axon branching (Figures 7*C-D,F,H-I*). However, we observed no difference in overall primary axon length measurements (Fig. 7*E*). Interestingly, there was also an increase in long collateral axon branches, including those longer 50μm, as compared to control neurons (Fig. 7*C-D, I*). Additionally, Dbn1-deficient neurons did not show an increase in axon protrusions, as seen following *Dbn1* OE *in vivo* (Fig. 1*G*, 2*B’’)*. Therefore, because Dbn1-deficient neurons form more axon branches but not more axon protrusions, *Dbn1* LOF regulates axon branch elongation *in vitro* in a manner similar to what we observe following LOF *in vivo* (Fig. 3-5).

To characterize *Dbn1* overexpression effects on cortical neuron morphology *in vitro,* we used a Dbn1 over-expression construct tagged with YFP to enable localization of Dbn1 in electroporated layer II/III neurons employing *ex vivo* electroporation (Geraldo et al., 2008). *Dbn1* OE in layer II/III CPNs cultured *in vitro* resulted in an increase in both axon branches and axon protrusions, compared to control neurons electroporated with GFP (Figs. 7*J-K’,M*). Further, there was no difference in primary axon length and average branch length in *Dbn1* OE neurons compared to control neurons *(data not shown*). Interestingly, in contrast to *Dbn1* LOF CPNs *in vitro*, there was no difference in the number of long axon branches (those greater that 50μm) compared to control neurons (Fig. 7*N*). These data suggest that *Dbn1* OE enables the formation of ectopic protrusions and short branches, but these branches fail to become long, stable, collateral branches, as seen in *Dbn1* LOF CPNs. The observation that *Dbn1* OE *in vitro* results in excess shorter branches could be due to the lack of extrinsic signals that are present in these *in vivo* conditions. These results are in line with our *in vivo* data showing that *Dbn1* OE results in an increase in axon branch initiation as assessed by protrusion number, but not branch elongation. Further, they highlight distinct phenotypes for *Dbn1* OE and LOF phenotypes, which can be assessed *in vitro* as well as *in vivo*.

### Dbn1 actin-binding and MT-coupling domains are necessary for Dbn1 overexpression-mediated protrusions

There are two major mammalian drebrin isoforms, E and A, and they are produced by alternative splicing from a single *Dbn1* transcript. Dbn1-E is encoded by the embryonic splice variant *Dbn1-E* and is expressed throughout the body. Within the brain, Dbn1-E is expressed prominently in the hippocampus and is also distributed across cortical layers within the developing somatosensory cortex (Ao et al., 2014). Dbn1-A, on the other hand, exhibits restricted expression in the brain, is enriched in dendritic spines and contains an additional amino acid sequence called “Insert 2” (Ins) (Koganezawa et al., 2017; Kojima, 2017; Shirao and Sekino, 2017). These Dbn1 isoforms contain several shared protein domains that include locations where Dbn1 binding partners associate and thereby facilitate cytoskeletal regulatory functions (Dun et al., 2012; Worth et al., 2013; Kojima, 2017; Shirao and Sekino, 2017). Dbn1 contains several functional domains, including: in its N-terminal region (eliminated below in the *Dbn1ΔN* deletion construct) an actin-depolymerizing factor homology domain (ADF-H), two actin binding domains (AB1/2) that include a coiled-coiled domain (CC) and helical domain (Hel); and in its C-terminal domain (eliminated below in the *Dbn1ΔC* deletion construct) a proline rich domain (PP), and a region implicated in MT plus-end coupling through End Binding Protein 1 (EB1) (Fig. 8*A*) (Shan et al., 2021). These domains associate with unique Dbn1 binding partners and serve specific cytoskeletal regulatory functions (Dun et al., 2012; Worth et al., 2013; Kojima, 2017; Shirao and Sekino, 2017).

**Figure 8.**
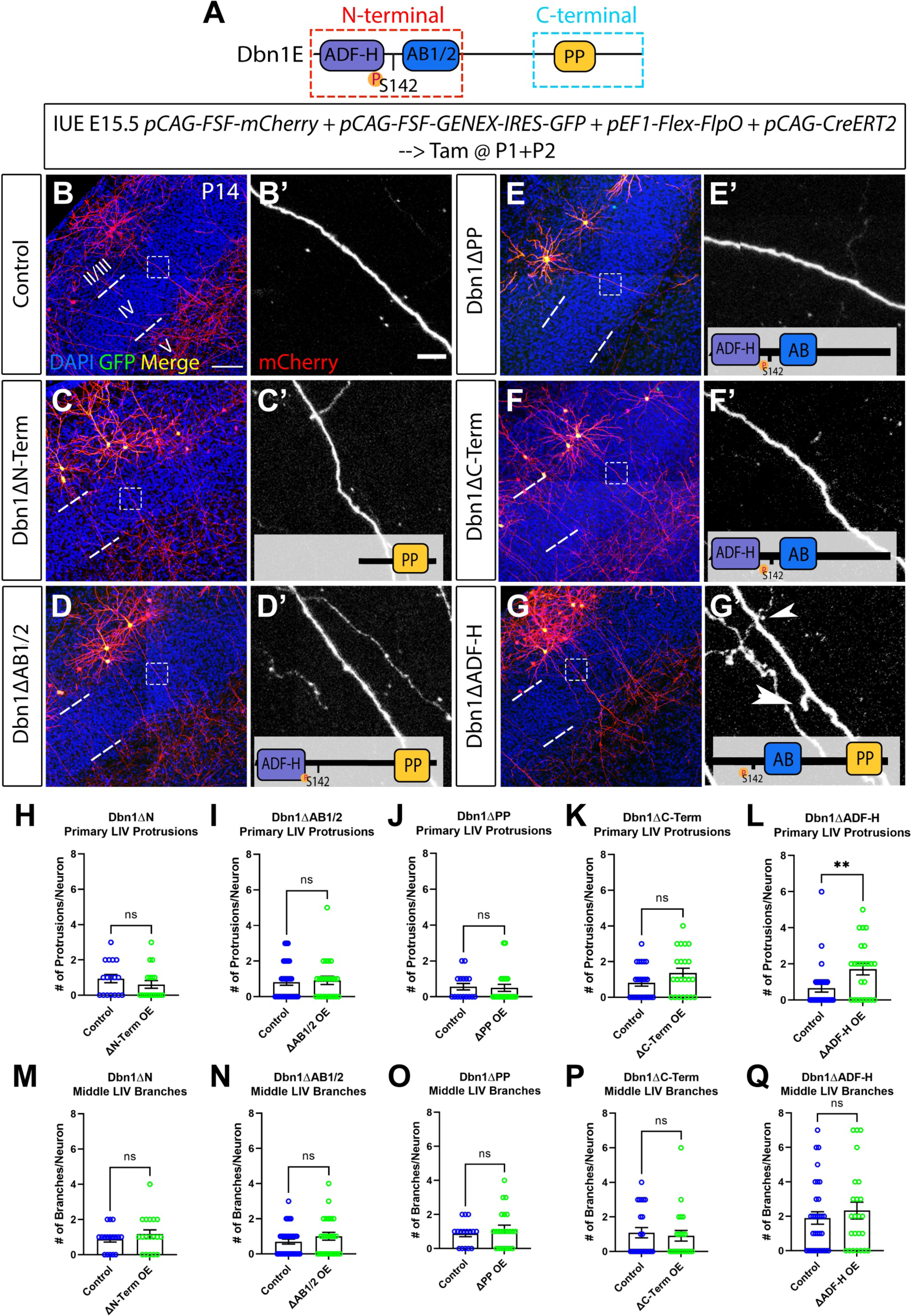
Actin-binding and MT-coupling domains of Dbn1 are necessary for protrusion formation upon overexpression. ***A,*** Diagram of Drebrin protein domain organization in Dbn1-E. ADF-H: actin-depolymerizing factor homology domain; S142: Cdk5 phosphorylation site; AB1/2: Actin Binding 1 and 2 domains; PP: proline-rich domain. ***B-G,*** Confocal immunofluorescent images of an electroporated layer II/III neuron within somatosensory cortical sections from CD1 mice that underwent electroporation at E15.5 via the bipartite approach with *pCAG-FSF-mCherry* (1ug/ul), *pEF1-Flex-FlpO* (500ng/ul), and *pCAG-CreERT2* (500ng/ul) and a control or specific deletion construct for each experiment detailed below. Tamoxifen was administered to electroporated pups at P1 and P2. Images of layer II/III neurons are stained with antibodies against dsRED and DAPI. ***B,*** *pCAG-FSF-GFP (2ug/ul).* Scale bars represent 100μm. ***C,*** *pCAG-FSF-Dbn1ΔN-terminal-IRES-GFP* (2ug/ul). ***D,*** *pCAG-FSF-Dbn1ΔAB1/2-IRES-GFP* (2ug/ul). ***E,*** *pCAG-FSF-Dbn1ΔPP-IRES-GFP* (2ug/ul). ***F,*** *Dbn1ΔC-terminal-IRES-GFP* (2ug/ul). ***G,*** *pCAG-FSF-Dbn1ΔADF-H-IRES-GFP* (2ug/ul). ***,B’,C’,D’,E’F’,G’*** mCherry labeling of zoomed in region of layer II/III axon within layer IV. Scale bar represents 25μm (B’). ***H-Q,*** For all experimental manipulations, only the layer IV region was traced for axon branches and protrusions since this was the region of interest for us. 18 *Dbn1ΔN-terminal* OE and 17 littermate control layer II/III neurons across 3 independent experiments with 2-3 pups per experiment were analyzed; 24 *Dbn1ΔAB1/2* OE and 33 littermate control layer II/III neurons across 2 independent experiments with 2-3 pups per experiment were analyzed; 22 *Dbn1ΔPP* OE and 16 littermate control layer II/III neurons across 1 experiment with 2-3 pups per experiment were analyzed; 22 *Dbn1ΔC-terminal* OE and 23 littermate control layer II/III neurons across 1 experiments with 2-3 pups per experiment were analyzed. 24 *Dbn1ΔADF-H* OE and 32 littermate control layer II/III neurons across 2 independent experiments with 2-3 pups per experiment were analyzed. Statistics: unpaired t-test. Lines represent mean and SEM. **p<0.01.

To investigate molecular mechanisms by which Dbn1 regulates collateral branch initiation and elongation, we generated *Dbn1E* overexpression constructs lacking select protein domains and assessed collateral axon branching and protrusion phenotypes within layer IV in CD1 WT mice *in vivo* using the BP approach. We focused on Dbn1E for this analysis since it is highly expressed during stages of cortical developmental when collateral branching events are initiated, whereas Dbn1A is expressed in later postnatal cortical development and spine maturation (Aoki et al., 2005). We observed that OE of *Dbn1ΔN* in layer II/III CPNs did not cause ectopic branching or protrusion phenotypes (Fig. 8*B-C’,H,M*). Similarly, OE of *Dbn1ΔAB1/2* in layer II/III CPNs did not cause any ectopic branching or protrusion phenotypes (Fig. 8*D,D’,I,N*), and neither did OE using *Dbn1ΔC* or *Dbn 1ΔPP*; this suggests that these domains are all necessary for Dbn1 OE axon protrusion phenotypes (Fig. 8*E-F’,J-K,O-P*). Interestingly, *Dbn1ΔADF-H* OE phenocopies full-length *Dbn1-E* OE, leading to axon protrusions within layer IV (Fig. 8*G-G’, L,Q*). Therefore, the ADF-H domain is dispensable for axon branch initiation as evidenced by the presence of protrusions in its absence (Fig. 7*8’,L*). We also assessed *Dbn1A* OE and observed no ectopic protrusion phenotypes, suggesting that Dbn1A and E play distinct roles in cortical development (*data not shown*).

Taken together, these results suggest diverse roles for Dbn1 domains in cortical layer II/III neuronal development. Each of the Dbn1 domains known to mediate F-actin bundling and MT-coupling is required for the formation of primary axon protrusions, which likely serve as the precursors for collateral axon branches (Hand et al., 2015; Ketschek et al., 2016; Shan et al., 2021).

### Phosphorylation of Dbn1 at S142 regulates interstitial axon protrusions

Dbn1 is phosphorylated by the Cdk5 serine/threonine kinase on serine 142 (S142), which lies within the Dbn1 N-terminal domain, and this posttranslational modification has been implicated in neuritogenesis, cell migration, growth cone turning, and synapse formation (Worth et al., 2013; Tanabe et al., 2014; Gordon-Weeks, 2017; Chen et al., 2022). Previous work identifies a role for Dbn1 phosphorylation by CDK5 at S142 in the regulation of F-actin bunding, which serves as a precursor for axon protrusions that in turn precede axon branch initiation (Worth et al., 2013; Kalil and Dent, 2014; Hand et al., 2015). To investigate further the molecular mechanisms underlying Dbn1 regulation of layer II/III neuron axon branching, we used DNA constructs encoding phosopho-mimetic (S142D) or phospho-dead (S142A) Dbn1 fused to a C-terminal YFP-tag in our BP electroporation screening paradigm and performed IUE at E15.5. *Dbn1^S142A^* over-expression did not cause any changes in overall layer II/III axon branching, including in the middle region of layer IV, or in protrusions across cortical layers compared to control neurons (Fig. 9*A-B’’,F,G*). Though we did not observe an overall increase in the total number of axon branches within the middle region of layer IV following *Dbn1^S142D^*over-expression, there was an increase in the number of total primary protrusions, similar to what we observed with WT *Dbn1* over-expression (Fig. 9*C-E*; Fig. 2*B-B’’*). These *in vivo* data suggest that phosphorylation of Dbn1 at S142 positively regulates axon protrusion formation since we do not observe ectopic protrusions when Dbn1 cannot be phosphorylated *(Dbn1^S142A^*over-expression) but do observe these phenotypes in the presence of constitutive Dbn1 phosphorylation (*Dbn1^S142D^* over-expression). Interestingly, *in vitro* analysis of cultured layer II/III cortical neurons shows that over-expression of both Dbn1 phospho-mimetic and phospho-dead mutants does not cause an increase in long axon branches (those greater than 50μm), as we observed in Dbn1-deficient neurons *in vivo,* but did cause an increase in axon protrusions, showing that both Dbn1 phospho-mutants mimic Dbn1 over-expression phenotypes rather than *Dbn1* LOF (*data not shown*). Taken together, these data show that Dbn1 OE regulates axon protrusion initiation in a manner dependent on the Dbn1 phosphorylation at S142.

**Figure 9.**
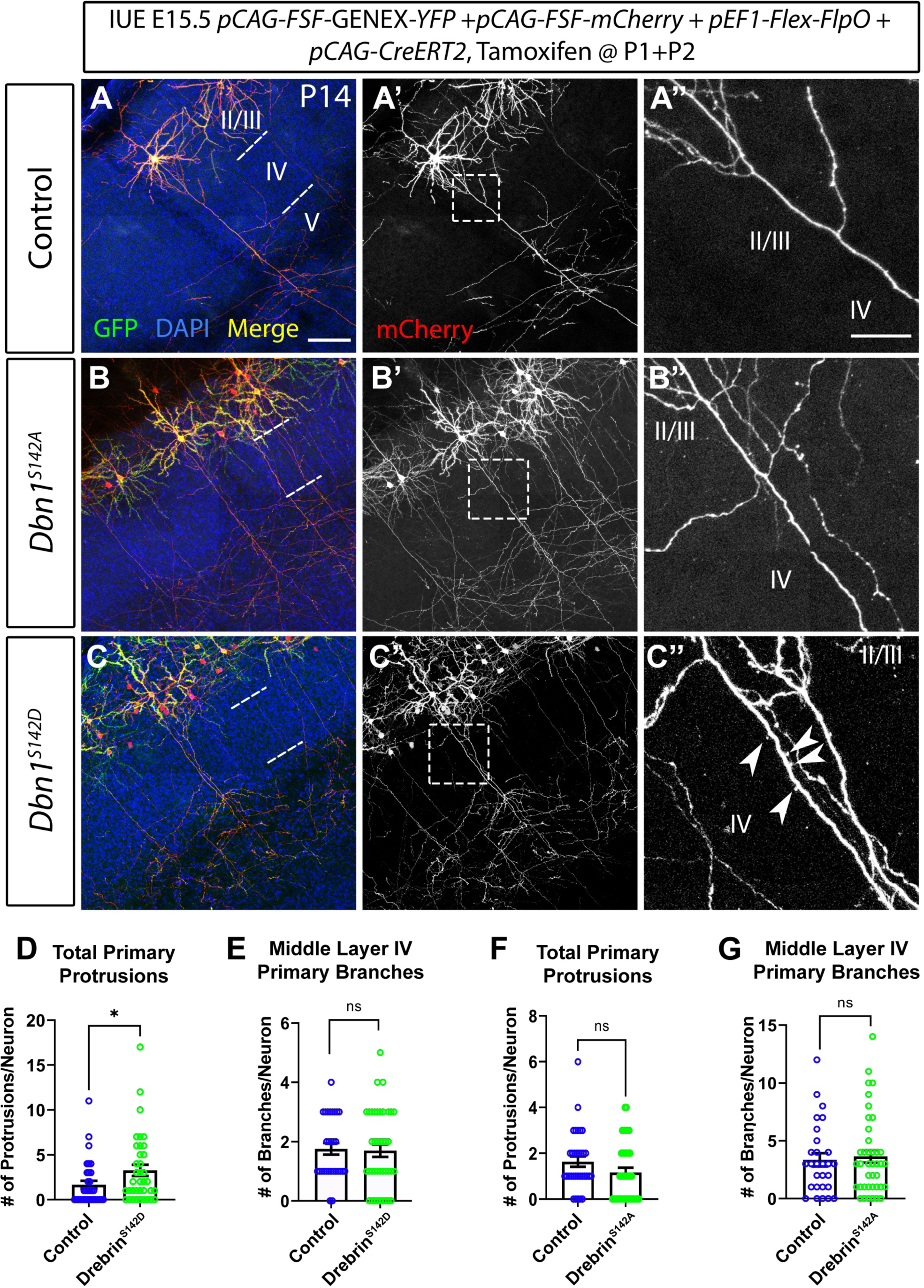
Phosphorylation of Drebrin at S142 regulates *Dbn1* over-expression axon protrusion initiation phenotype. ***A-C’’,*** Confocal immunofluorescent images of P14 layer II/III neurons from CD1 following IUE at E15.5 using the bipartite approach. Sections were stained with antibodies against dsRED, against GFP to detect YFP, and DAPI. Neurons were electroporated with the following constructs: *pCAG-FSF-mCherry* (1ug/ul), *pEF1-Flex-FlpO* (500ng/ul), *pCAG-CreERT2* (500ug/ul), and individual plasmids for each experimental manipulation. Tamoxifen was administered at P1 and P2. ***A,*** Merged image of an example control layer II/III neuron electroporated with *pCAG-FSF-GFP* (2ug/ul) in addition to bipartite plasmids. Cortical layer boundaries are demarcated using DAPI staining. Scale bar represents 100μm. ***A’,*** mCherry labeling of control neuron. Boxed region is magnified in *A’’*. ***A’’,*** Zoomed in image of region of layer IV along layer II/III axon. No atypical branching or protrusion phenotypes are observed. Scale bar represents 50μm. ***B,*** Example of a layer II/III neuron over-expressing the phospho-mutant Dbn1 mutant construct *pCAG-FSF-Dbn1S142A-YFP* (2ug/ul). ***B’,*** mCherry labeling of layer II/III neuron. Boxed region is expanded in *B’’*. ***B’’,*** Zoomed in image of a region of axon within layer IV. No atypical morphologies are observed. ***C,*** Example of a layer II/III neuron over-expressing the phosphomimetic Dbn1 mutant construct *pCAG-FSF-Dbn1^S142D^-YFP* (2ug/ul) in addition to other listed bipartite constructs. ***C’,*** mCherry labeling of layer II/III neuron. Box region is amplified in *C’’*. ***C’’,*** Zoomed in image of layer IV region along layer II/III neuron axon. The presence of Axon protrusions in this region is observed. ***D-E,*** Quantification of layer II/III neurons over-expressing Dbn1^S142D^. 33 control neurons and 37 *Dbn1^S142D^* OE neurons were analyzed from 2 independent experiments across 2 mice per experiment. Statistics: unpaired t-test. Lines represent mean and SEM. *p<0.05. ***F-G,*** Quantification of layer II/III neurons over-expressing Dbn1^S142A^. 29 control neurons and 40 *Dbn1^S142A^* OE neurons across 2 independent experiments from 2-3 mice per experiment.

### Phosphorylation of Dbn1 at S142 is not required for suppression of collateral branching in layer IV

To determine whether phosphorylation of Dbn1 at S142 is necessary for the normal suppression of collateral axon branching by Dbn1, we performed rescue experiments using the conditional *Dbn1^flox^* allele and phosopho-mimetic or phospho-mutant constructs in a slightly modified labeling version of our BP approach (see Methods). As we saw previously, *Dbn1^S142D^*over-expression resulted in an increase of axon protrusions in *Dbn1^fl/+^* layer II/III neurons, but not in *Dbn1^fl/fl^* layer II/III neurons (Fig. 4*C-C’’*, *F’*). Further, *Dbn1^S142D^* expression in the context of *Dbn1^fl/fl^* LOF eliminated the ectopic axon branching phenotype observed in *Dbn1* LOF neurons (Fig. 4*C-C’’,G*). On the other hand, as expected, *Dbn1^S142A^* expression in *Dbn1^fl/+^* layer II/III CPNs did not result in an increase in axon protrusions within layer IV (Fig 4*D-D’’*,*E*). Interestingly, *Dbn1^S142A^* expression in layer II/III neurons suppressed the *Dbn1* LOF layer IV ectopic branching phenotype (Fig. 4*D-D’’,F*). These results suggest that Dbn1 normally suppresses ectopic collateral branching within layer IV and that phosphorylation at S142 is not required for this aspect of branch regulation.

## Discussion

We address here the molecular mechanisms underlying spatially regulated interstitial neocortical lamination axon branching using newly developed approaches to study candidate signaling molecules that regulate cytoskeletal dynamics. We show that Dbn1, a cytoskeletal regulatory protein that links microtubules and actin, regulates collateral axon branching of layer II/III cortical projection neurons (CPNs) *in vivo* and *in vitro*, playing a dual role in this process. *Dbn1* LOF results in layer II/III CPN collateral axon branch elongation and stabilization in layer IV, whereas *Dbn1* OE leads to axon branch initiation.

We have shown previously that filopodial protrusions initially form along layer II/III CPN axons in all cortical layers, however only those in target layers (II/III and V) are stabilized and elongate to form collateral branches (Hand et al., 2015). *Dbn1* OE shows that Dbn1 is sufficient for this initial interstitial protrusion formation step that serves as a precursor for collateral branch formation (Dent et al., 2004; Gibson and Ma, 2011; Kalil and Dent, 2014; Hand et al., 2015). *Dbn1* over-expression (OE) results in an increase in short axon protrusions along the length of the CPN primary axon, including layer IV. These ectopic protrusions are present early in postnatal development at P7 and remain at P14, when laminar CPN axon collateral branching is almost complete (Fig. 2, *data not shown*). Further, we show that Dbn1 phosphorylation at S142, and also Dbn1 protein domains known to mediate F-actin bundling and MT-coupling, are necessary for mediating OE axon protrusion phenotypes.

In addition, *Dbn1* LOF results in an increase in the number of layer II/III CPN collateral axon branches *in vivo*, in particular within the middle region of layer IV, and thus is necessary to inhibit the development of mature collateral branches in this region where they rarely occur. We further show with rescue experiments that Dbn1 is sufficient to restrict ectopic branching in layer IV, and that this is independent of S142 phosphorylation. *In vitro, Dbn1* LOF also results in collateral axon branch elongation, further supporting a role for Dbn1 in the regulation of collateral axon branching. We found that *Dbn1* LOF phenotypes *in vivo* were only observed following loss of Dbn1 expression within a developmental window between the completion of layer II/III radial cell migration (P0) and axon branch initiation (∼P3). *Dbn1* LOF at P4 or at P7, after the developmental time interval during which collateral branches are initiated, does not result in axon branching phenotypes.

Because *Dbn1* OE and LOF exhibit distinct phenotypes with respect to CPN protrusions and collateral axon branching, we performed a series of Dbn1 structure-function experiments to investigate the underlying molecular mechanisms. Our data show that the Dbn1 AB1/2 domains within the N-terminal region, and also the PP domain within the C-terminal region, underlie *Dbn1* OE production of axon protrusions: axon branch initiation events that precede the formation of mature collateral axon branches. The ADF-H domain, however, is dispensable for these Dbn1 OE protrusion phenotypes. This aligns with previous studies showing the Dbn1 ADF-H domain does not influence filopodia formation in COS-7 cells (Worth et al., 2013).

Dbn1 exists in a repressed conformational state, that is relieved by Cdk5 phosphorylation, such that an as yet undefined region in the C-terminus binds to a region at the border between the ADF-H and CC domains, both of which lie within the Dbn1 N-terminal region (Worth et al., 2013). In this conformation Dbn1 can bind to individual F-actin filaments but cannot form actin bundles, a necessary step for collateral axon branch initiation (Worth et al., 2013; Pacheco and Gallo, 2016). The S142 phosphorylation site lies between the ADF-H and CC domains and is therefore occluded in this closed conformation (Worth et al., 2013). However, Cdk5 phosphorylation of Dbn1 at this site relieves this intramolecular occlusion, enabling Dbn1 to bundle F-actin and interact with EB3 at the +TIPs of MTs, ultimately resulting in the initiation of collateral axon branches first observable by the formation of actin patches, which are similar to the protrusions we have previously shown to precede collateral axon branching and describe here (Worth et al., 2013; Ketschek et al., 2016; Pacheco and Gallo, 2016). We found that OE experiments using the phospho-mimetic mutant Dbn1^S142D^ caused ectopic protrusion formation along layer II/III CPN axons but not an increase mature collateral axon branch number, whereas overexpression of the non-phosphorylatable Dbn1^S142a^ did not cause either protrusion or axon branching phenotypes. Therefore, this mode of actin bundling enabled by S142 phosphorylation may be required for Dbn1 to initiate collateral branch formation in layer II/III CPN axons. In addition, previous work using bioinformatic modeling predicted that End Binding 1 (EB1), a microtubule plus-end protein, binds to Dbn1 in the C-terminal region downstream of the PP domain (Shan et al., 2021). Therefore, MT and actin binding properties of Dbn1 found within both the N and C-terminal regions are likely necessary for Dbn1 regulation of collateral branch initiation.

We observe in our rescue experiments that Dbn1 suppression of collateral axon branching in layer IV is independent of Dbn1 phosphorylation at S142. These results indicate that suppression of collateral branching is independent of actin bundling and may support a role for MT depolymerization in the suppression of collateral axon branching. Alternatively, suppression of collateral axon branch elongation in the absence of S142 phosphorylation may involve Dbn1 inhibition of actin bundling owing to its association with and sequestration of individual actin filaments; localized Cdk5 phosphorylation could serve to provide spatial regulation of this process. We observe that epitope-tagged Dbn1 and Dbn1 phosphomutants localize to the axon region within the middle of layer IV upon OE, showing that Dbn1 is indeed positioned to mediate the ectopic branching phenotypes we observe in these experiments. Interestingly, upon OE of the *Dbn1^S142D^* phosphomimetic in *Dbn1^fl/fl^*mutant CPNs we do not observe the induction of ectopic axon protrusions, suggesting that in order to induce axon protrusions, Dbn1 levels must be very high and that we were likely unable to achieve sufficient expression to induce protrusions in Dbn1-deficient layer II/III CPNs.

Our observations provide an initial example of a molecule that regulates interstitial axon collateral branching *in vivo* in a population of CPNs that normally forms laminar-specific axon branching patterns. They are in agreement with previous findings showing that Dbn1 over-expression in sensory neurons causes an increase in filopodial-like protrusions (Dun et al., 2012; Ketschek et al., 2016). However, our results are in contrast to other results showing that *Dbn1* LOF results in reduced axon branching in sensory neurons (Ketschek et al., 2016). This discrepancy may reflect inherent differences between CNS and sensory neurons. In addition, our data complement work demonstrating roles for intracellular signaling components, including WNK kinase and AnkB440, in the maintenance and refinement of terminal axon branches and contralateral colossal axon projections, respectively (Creighton et al., 2021; Izadifar et al., 2021).

The precise patterning of layer II/III CPN collateral branching will determine synaptic connectivity patterns these neurons ultimately adopt (Sanes and Yamagata, 1999, 2009; Sanes and Zipursky, 2020; Dorskind and Kolodkin, 2021). Therefore, extending ectopic branches in regions where collateral branching normally does not occur has the potential to alter overall circuit formation and neuronal wiring. Our results suggest that Dbn1 must be inhibited to achieve the normal maturation pattern of interstitial laminar-specific axon branch elaboration. We hypothesize that branch promoting cues are present in layers II/III and V, where layer II/III CPN collateral branches are typically stabilized, and that inhibitory cues are present in layers IV and VI, where collateral branching seldom occurs (Fig. 10*A*). Therefore, branch promoting cues may inhibit Dbn1 to enable elongation of collateral branching in these target regions, while the absence of these cues in layer IV may prevent the formation of stable branches (Fig. 10*B,C*). It will be interested to investigate whether or not these cues act through Cdk5 to activate axon branch initiation. Future studies will address the functional and behavioral implications of *Dbn1* OE and LOF phenotypes.

**Figure 10.**
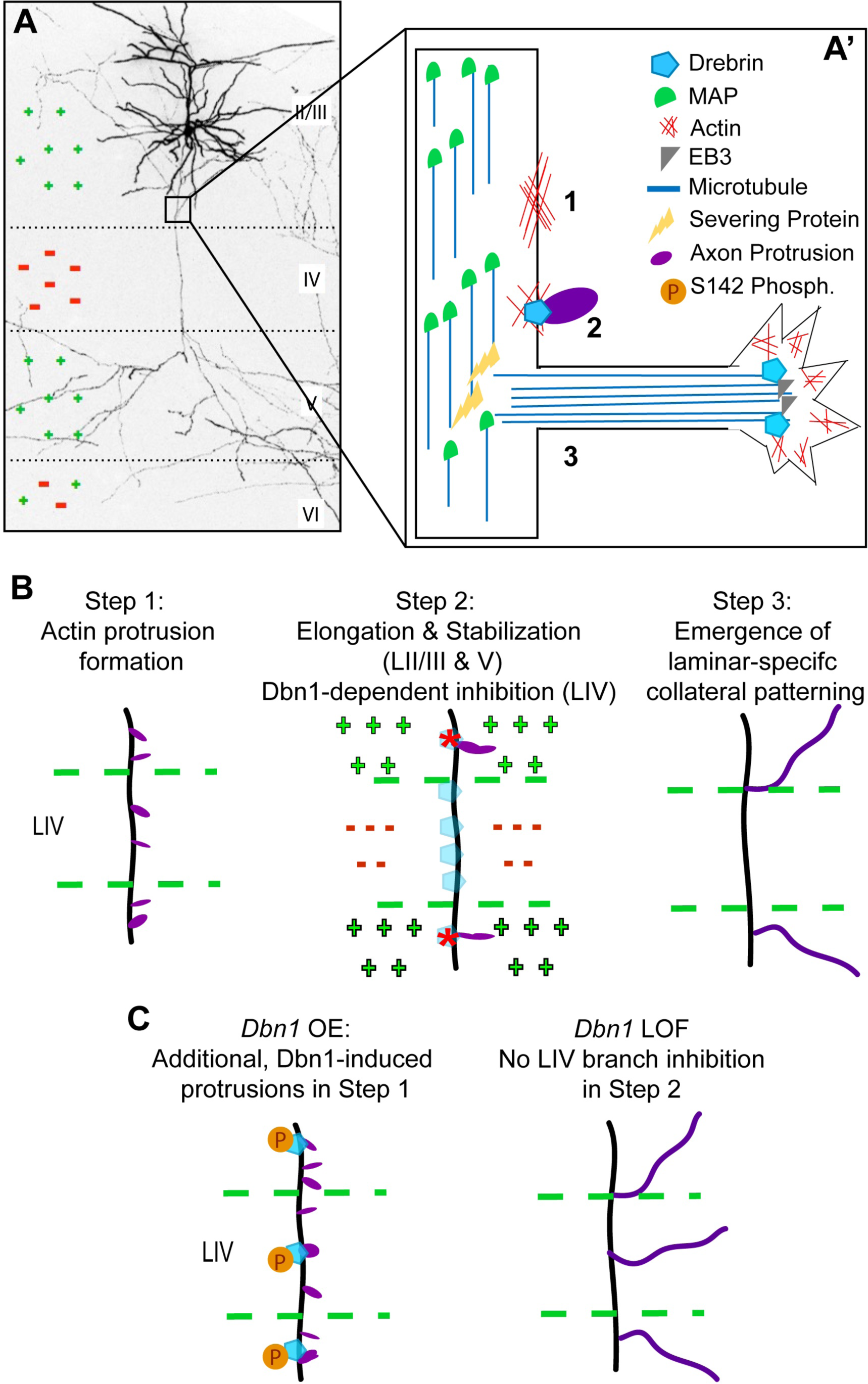
Model for Drebrin-mediated regulation of axon branching and protrusion in layer II/III neurons. ***A,*** Confocal maximum intensity image of a layer II/III neuron electroporated with pCAG-LSL-GPF (2ug/ul) from P14 somatosensory cortical sections. Layer II/III neurons typically branch in layer II/III and V and branches are usually absent from layer IV. Positive extrinsic cues likely enable collateral branch extension in layers II/III and V (green plus signs) and negative extrinsic cues in layer IV likely inhibit collateral branch extension in this region (red minus signs); a mixture of cues in layer VI is also possible. ***A’,*** Depiction of microtubule associated proteins (MAPs) within a collateral branch on layer II/III neurons. Microtubules within the axon shaft are stabilized by MAPs. Severing proteins, such as Spastin and Katanin, destabilize microtubules, enabling them to invade actin precursor meshwork and elongate into a stabilized collateral branch. Drebrin interacts with both actin and microtubules and may regulate collateral branch initiation and/or elongation through these interactions. ***B,*** Role of Drebrin in the initiation and suppression of collateral axon branches. Step 1: Actin protrusions are formed along the length of the layer II/III CPN axon. Step 2: These protrusions are then stabilized and form elongating collateral branches in target layers (II/III and V) and are retracted so do not form branches in layer IV. We show that this suppression is mediated by Dbn1. We hypothesize that there are branch-promoting cues in layers II/III and V that enable this elongation and branch inhibitory cues, or Dbn1 activating cues, in layer IV that prevent elongation in this region. Step 3: Subsequent regulation of axon branch development results in the emergence of stereotypic laminar-specific collateral axon branch patterns. ***C,*** *Dbn1* OE and LOF phenotypes. *Dbn1* OE leads to ectopic, stable, axon protrusions mediated by phosphorylation at S142, while *Dbn1* LOF leads to ectopic, stable, collateral branch formation in layer IV.

Since Dbn1 is an actin/MT binding protein that is ubiquitously expressed in the neocortex, it will be important investigate how extrinsic cues regulate the laminar-specific innervation branching phenotypes we observe in the *Dbn1* mutant CPNs. Dbn1 acts in concert with afadin to stabilize the cell adhesion protein nectin at endothelial junctions (Rehm et al., 2013), and it also regulates T cell activation by interacting with CXCR4 and actin in CD4^+^T cells (Perez-Martinez et al., 2010). These previous studies highlight the potential for extrinsic cues to regulate Dbn1, and identification of these cues in the context of cortical lamination is necessary to further elucidate molecular mechanisms underlying neocortical circuit formation (Perez-Martinez et al., 2010; Rehm et al., 2013). The genetic tools generated this study, in combination with our observations relating to Dbn1 regulation of axon collateral branch formation, lay the groundwork for investigating the extrinsic cues that define laminar-specific axon branch innervation in layer II/III CPNs and, potentially, in other populations of projection neurons the mammalian neocortex.

## Acknowledgements

We thank members of the Kolodkin laboratory for thoughtful input on this work and Nicole Kropkowski for maintaining mouse colonies used in this study. We thank Natalie Hamilton for help with statistical analyses and calculations, and Yatindra Awasthi for his assistance. We also thank Ulrich Müller and Soraia Barao for guidance and collaboration, and Jeremy Nathans and Feng-Quan Zhou for guidance and sharing reagents. We thank Noah DeMarco, MS, for coding the two python programs that were essential to the completion of this study. This work was supported in part by the Howard Hughes Medical Institute (ALK), the CMM Graduate Training Program at The Johns Hopkins University School of Medicine–T32-GM008752 (JMD), the Kavli Neuroscience Discovery Institute at Johns Hopkins (SS), and an EMBO Postdoctoral Fellowship no. 364-2021 (JZ).

## Notes

### Competing Interest Statement

The authors have declared no competing interest.

